# Validation-free estimation of chronological age via close-kin

**DOI:** 10.64898/2026.06.01.721774

**Authors:** Luke R. Lloyd-Jones, Mark V. Bravington, Hien D. Nguyen, Robin Thomson, Jack H. Easton

## Abstract

Age is a fundamental life-history parameter in animal ecology and wildlife management. Age informs key ecological characteristics including population age structure, recruitment strength, extinction risk, reproductive maturity, and mortality rates. This importance has necessitated the development of chronological age estimation methods for wild animals. However, estimating chronological age is challenging for wild species, with noisy and potentially biased measures typically gathered from morphometrics, physical characteristics or, more recently, molecular methods like DNA methylation. These measures of age require at least some initial validation set of known-age individuals, or known time intervals, which is difficult to obtain for many species. Here, we present a solution to inferring the relationship between chronological age and error-prone observed age that does not require known-age individuals. The model couples the formulae for occurrence rates of half-sibling pairs, which decrease as a function of the birth-year gap between two sampled individuals, with time of capture. A pseudo-likelihood framework is developed for parameter estimation that can resolve linear and non-linear relationships and provide variance parameter estimates. We explore the method’s efficacy for estimating chronological age using forward-in-time simulation and validate prior estimates of the relationship between vertebral band counts and chronological age for 3,000 school shark (*Galeorhinus galeus*) from an Australian fishery.

## 1. Introduction

Chronological age is a fundamental life-history parameter in animal biology, population dynamics, and ecology. In wildlife management, age informs the understanding of key ecological characteristics such as population age structure, recruitment strength, extinction risk, reproductive maturity, and mortality rates (Jennings et al., 1998; Campana, 2001; Nussey et al., 2008; Sherley et al., 2014). The importance of chronological age in animal ecology has necessitated the development of age estimation methods for wild animals.

For many wild species, physical measurements like counting growth bands in hard parts e.g., otoliths, vertebrae, teeth or fin spines (Jones, 1971; Francis and Campana, 2004; Walker et al., 2001) or using bone age determination (Schucht et al., 2021), can provide accurate age measurements but typically require dead specimens. For endangered species, lethal methods are not an option, leading researchers to explore morphometrics and teeth wear (Todd et al., 2018). Inference of age from length using growth curves is possible but has limitations including age-at-length variance across populations and individuals. Furthermore, bias from length measurement error and selective sampling need to be properly accounted for to age individuals (Gwinn et al., 2010). Growth typically slows or ceases at maturity, reducing the utility of age-at-length estimation, and impeding hard part measurement through band narrowing (Harry, 2018; Natanson et al., 2018). Epigenetic clock research is a rapidly advancing and promising avenue with a wide range of wild species being studied and many accurate estimates of age generated (Polanowski et al., 2014; Stubbs et al., 2017; Thompson et al., 2017; Jarman et al., 2015; Lu et al., 2023). However, these epigenetic studies use an initial calibration data set with both known chronological age or low-noise age scores — a measurable quantity related on average to chronological age, such as hard part measures, body length, or DNA methylation levels — which can be very challenging to obtain for many species.

Mark-recapture methods have been used to estimate age non-lethally for individuals within the study (Eveson et al., 2004). For example, repeat photo identification via distinctive markings was used for humpback whales and coupled with DNA methylation to estimate age (Polanowski et al., 2014), and recapture identification after banding as chicks for short-tailed shearwaters (De Paoli-Iseppi et al., 2019). For species where marking and recapture is challenging, indirect genetic methods have been used to identify repeat samples (Luikart et al., 2010), with studies collecting passive repeat samples from faeces or hair (Kendall et al., 2008).

The close-kin mark-recapture (CKMR) method leverages genotyping technology to identify genetic tags in the form of shared DNA between closely related individuals (Skaug, 2001; Bravington et al., 2016). CKMR can estimate total adult population size, rate of change, and survival rates, overcoming many traditional challenges of surveying cryptic or elusive species. The most commonly used kinship types in CKMR are parent-offspring pairs (POPs) and half-sibling pairs (HSPs), i.e., individuals sharing exactly one parent. CKMR has been successfully applied to aquatic species, such as southern bluefin tuna (Bravington et al., 2016), salmon (Wacker et al., 2021), white, school, grey nurse, and northern river sharks (Hillary et al., 2018; Bradford et al., 2018; Bravington et al., 2019; Thomson et al., 2020), and to a handful of terrestrial species (Lloyd-Jones et al., 2023). Parameter estimation from CKMR data relies on kinship probability formulae for pairs of sampled individuals. Those formulae depend on the dates-of-birth of the samples, since that is the moment at which offspring “mark” their parents. Birth-date is not usually measurable directly but must be inferred from date and age when sampled, so it is important to have at least some information about age. The noisier that information is, the less precise will be the demographic parameter estimates. If age estimates are biased, or if their uncertainty is ignored, there will be bias in parameter estimates too (Petersma et al., 2024). While age-free analyses are theoretically possible, practical applications have yet to demonstrate their viability.

We develop a new framework for estimating chronological age using multi-year HSPs. We were motivated by the need for validation-free methods for chronological age estimation and to calibrate existing methods of indirect age estimation (hereafter referred to as auto-calibration). Furthermore, we intended to integrate the method with emerging DNA methylation-based age estimators, which when coupled with minimally invasive sampling, a single tissue biopsy could yield both close-kin information and age. The solution rests on the notion that pairwise rates of HSPs decrease as birth-date gap increases because each parent of the first born has more opportunity to die as the interval widens. In a multi-year CKMR study, this will affect how the empirical rate of HSP-finding varies with (i) increasing gap between sample years and (ii) increasing gap between age-score measurements. Both are driven by the same phenomenon, so auto-calibration amounts conceptually to finding a scaling for the age score at which the two rates coincide, perhaps also allowing for noise in the age score. We explain this concept further in Section 2.1, then introduce a basic CKMR HSP setup in Section 2.2, which we use to illustrate how auto-calibration fits into a CKMR model. Moving from the conceptual to the practical, Sections 2.3 and 2.4 present two statistical approaches for auto-calibration: a simple one that assumes a linear relationship between age score and chronological age but neglects noise in the age score (Section 2.3); and a more complex one that can allow for non-linearity and noise (Section 2.4). We test the method using forward-in-time simulation in Section 3. In Section 4, the more-complex model is applied to the calibration of vertebral band count for school shark (*Galeorhinus galeus*) from 3,000 samples taken for a CKMR study from a shark fishery in southern Australia. Ageing by vertebral band count is used here as one instance of a broader class of noisy age measurements, including DNA methylation-based estimates.

## 2. Methods

### 2.1 Conceptual basis for auto-calibration

The data consist of an age score *G*_*i*_ for each individual *i* ∈ {1, …, *n*} on an arbitrary scale. This score could be a composite, e.g., from a factor analysis over many DNA methylation sites, or a single strongly associated measure, such as length, or band counts from otoliths or vertebrae. For each individual, we assume that we also have genotype data that are adequate for reliably inferring kinship between pairs of individuals.

Our solution uses HSPs to calibrate the relationship between chronological age and the observed age score. Considering maternal half-siblings for illustration, the probability that any two sampled individuals are an HSP decreases with increasing interval between the individuals’ birth dates, assuming constant adult fecundity. We can use kinship information and this property of decreasing HSP rates at greater birth-gaps to estimate individual chronological age from the noisy age score. Below we assume that *G*_*i*_ is a linear function of chronological age and later extend this to other potential functions. Conceptually, this can be achieved by:

1. Look at all pairwise comparisons of sampled individuals (*i, j*) that were caught in the same year. The difference in their age score, Δ*G*_*ij*_, will linearly reflect the difference in their birth years. The HSP rate should decrease exponentially with increasing Δ*G*_*ij*_. Under the standard demographic assumption of a constant instantaneous mortality rate, across adult ages the probability of surviving an interval is modelled as an exponential decline with birth-gap.
2. Investigate all pairs regardless of year sampled, where Δ*G*_*ij*_ is near 0. These individuals were caught at similar ages but in different years. As the difference in sample years increases, the rate of HSPs should also decline exponentially.
3. Finally, find the scaling of *G* that makes these two rates the same. The ratio of the two rates of exponential decline converts the noisy score *G*_*i*_ into individual age (*A*_*i*_).

This conceptualisation ignores a large amount of information because the great majority of pairwise comparisons will be unused. Sections 2.3 and 2.4 present two statistical approaches that formalise and extend this concept to include all comparisons: a simple approach that assumes negligible noise in the age score (Section 2.3), and a fully marginalised approach that accounts for noise explicitly (Section 2.4). Both build on a common CKMR HSP model introduced in Section 2.2.

### 2.2. Simple half-sibling CKMR model

Let *b* be birth year, *a* be realised age of the random variable *A, G* be a random variable with realisation *g*, and *y* be observed year of sampling. Subscripts *i* and *j* index over individuals. We consider a simple CKMR setting where abundance is stable over time, mortality rate is constant across adult ages and over time, breeding is annual, and fecundity does not change, on average, with age. Bravington et al. (2016) detail the HSP probability calculation for CKMR; here we present a simplified form sufficient for auto-calibration.

For a pair of individuals with birth years *b*_*i*_ and *b*_*j*_, the probability that they are a maternal HSP (*K*_*ij*_ = MHSP) depends on birth-gap as follows:

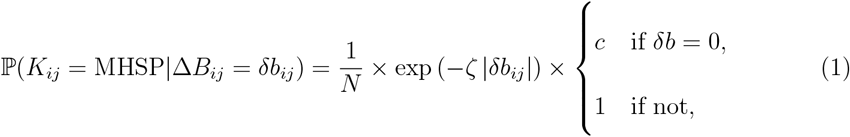

where *ζ* is the adult mortality rate, *N* is the female adult abundance parameter assumed constant through time, and *c* adjusts the within-cohort HSP rate, which can differ relative to the baseline model due to, e.g., correlated survival among littermates with multiple paternity or conversely for species that only produce one child per breeding event. We denote the demographic model parameters ***θ*** = (*N, ζ, c*)^t^. The absolute value |*δb*| enters (1) as we do not know the true birthdate of the compared individuals and therefore do not know which is older. Equation (1) is sufficient for illustration, but in other settings HSP probability formulae can be more complicated; Bravington et al. (2016) explain how to generalise the principles.

Each pairwise comparison of sampled individuals assessed for kinship can be modelled as a Bernoulli random variable with success probability stipulated by equation (1). Equation is adequate for a sex partitioned model but for aggregated sexes the probability that the pair share either a mother or a father requires a factor of four (see Methods of Hillary et al. (2018)). The estimation proceeds by maximising the pseudo-log-likelihood

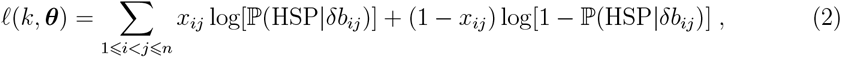

where *x*_*ij*_ is an observed binary indicator for whether individuals *i* and *j* are a match for an HSP generated from the genetic data. Equation (2) is a pseudo-likelihood because the pairwise comparisons are not strictly statistically independent e.g., because no animal can have more than one mother. However, pseudo-maximum-likelihood estimates are asymptotically unbiased, and variance estimates are accurate when sample sizes are small relative to the size of the population (see Section 4 of Bravington et al. (2016) for a discussion and Supplement of McDowell et al. (2022)).

### 2.3 Low-noise approach: pairwise differences

We assume for now that our noisy age measure *G* is unbiased with constant variance, so that

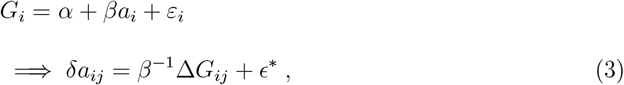

where ϵ^*^ = *β*^−1^ (*ε*_*j*_ − *ε*_*i*_) and Var(*ε*_*i*_) = *σ*^2^. For each individual, we do not observe *a* or their birthdate *b* directly so instead we use the fact that *b*_*i*_ = *y*_*i*_ − *a*_*i*_.

If the noise in the age score is small relative to the slope of the chronological age to age score relationship, the latent ages can be treated as approximately known. Using *b*_*i*_ = *y*_*i*_ − *a*_*i*_ and ignoring the error term in equation (3), the birth-gap can be expressed in terms of observed quantities:

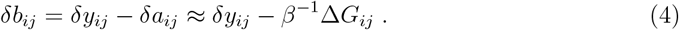

Substituting into equation (1) gives the HSP probability in terms of observed quantities:

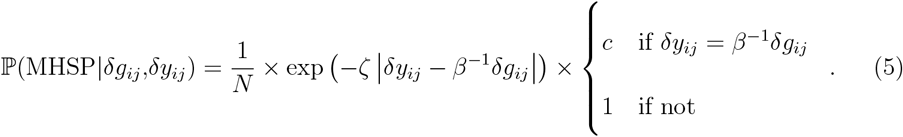

Each pairwise comparison for kinship is modelled as a Bernoulli random variable with success probability given by equation (5). We include *c* for each pair when *δy*_*ij*_ − *β*^−1^*δg*_*ij*_ = 0 up to a small tolerance, appropriate when the age-score variance relative to the slope is small. We denote the chronological age to age score relationship parameters ***η*** = (*α, β, σ*^2^)^t^, where *σ*^2^ represents the conditional variance of the age score given chronological age, which is assumed negligible and not estimated. The parameter *α* is further not recovered from the estimation process; the absolute scale of the estimated relationship requires extra information to be determined.

This formulation has the advantage of simplicity: auto-calibration reduces to estimating the scaling *β* that aligns the two rates of HSP decline described in Section 2.1. However, as measurement noise increases, the approximation degrades. In Section 2.4 we present a generalisation that accounts for the chronological age to age score relationship noise explicitly by marginalising over latent ages. Supplementary Note **??** shows that the parameters (*N, ζ, β, c*) of the pairwise difference model are identifiable: four distinct contrast points in (*δy, δg*)-space suffice to uniquely determine all four parameters.

### 2.4 Full-marginalisation approach

In many studies the variance in *G*_*i*_ given chronological age may not be small and should be modelled adequately. Rather than treating latent ages as approximately known, we marginalise over all possible chronological ages for each individual.

Suppose we observe realisations *g*_1_ and *g*_2_ for two individuals sampled in *y*_1_ and *y*_2_ respectively, then we need to evaluate the following expression for their HSP probability from the law of total probability:

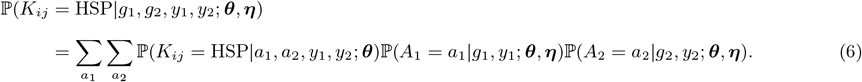

In equation (6), *a*_1_ and *a*_2_ span the plausible support determined by the distribution for *A* conditioned on *g* and *y*. Equation (6) makes explicit a marginalisation over the latent ages of both individuals. In the limiting case where ℙ (*A* = *a*|*g, y*) concentrates at a single age, the double sum in (6) collapses to evaluation at those point estimates, recovering the pairwise difference approximation. Supplementary Note **??** shows this formally.

Implicitly in equation (6), we condition on the fact that the animals were actually sampled (*S*), i.e., they may not be equiprobable draws from the population, but instead may be subject to some selectivity, e.g., through size selectivity by fishing gear.

To evaluate equation (6), we apply Bayes’ rule:

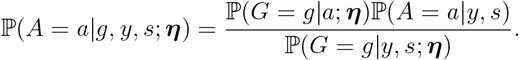

From the data, we have the empirical marginal distribution 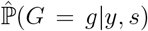, which we can model as

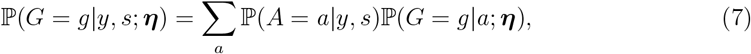

and we incorporate a procedure to estimate a smooth model for ℙ (*A* = *a*|*y, s*), representing the age distribution of sampled individuals conditional on capture year, which is updated jointly within the overall fitting algorithm. Supplementary Notes **??** and **??** show how we use discrete deconvolution via smoothing to incorporate equation (7) into the likelihood procedure. We refer to this as the “full-marginalisation” method. We note that in other settings where selectivity is modelled explicitly (e.g., fish stock assessment (Maunder and Punt, 2013)) or can be assumed uniform, deconvolution may not be required. Supplementary Notes **??** and **??** show the necessity of some known-age samples to estimate *α* and the identifiability of the parameters of the full-marginalisation model under an assumption of a stable age distribution for the chronological age prior.

In the above description, the CKMR demographic probabilities depend only on ***θ*** once individual ages are fixed. Age-measurement parameters ***η*** affect the posterior distribution of latent ages, which in turn propagates uncertainty into the kinship likelihood through the marginalisation over age.

## 3. Simulation study

### 3.1 Simulation design

To investigate the performance of the method and validate the implementation, we used the `CKMRpop` (Anderson, 2022) software to perform forward-in-time simulations of age-structured populations in the R programming language (R Core Team, 2021). `CKMRpop` is highly flexible in allowing for the parameterisation of demographic features such as individual birth and death cohorts, age and sex-specific annual census sizes, identities and ages of sampled individuals, and tracks the complete pedigree of all individuals over the simulation period. Importantly, `CKMRpop` features a fast method to recursively traverse the simulated pedigree and identify all pairwise relationships between the sampled individuals.

Our simulation was loosely based on some, but not all, of the demography of school sharks presented in Section 4. We simulated a stable population size over 100 years. We sampled at years 90–97, when there were ≈ 13,000–13,500 individuals in the population (approximately 1,000 adults), which balanced population size and computation time. The age at maturity was set at 11 years. Fecundity was assumed constant after maturity for males and females. Survival for the females and males was an ogive across the age range with minimum value of 0.75 and maximum of 0.82; adults varied from 0.79 to 0.82 with an average across all adults of 0.805 computed from the observed simulated data across simulation scenarios. We set the maximum age at 25, after which all animals died; this had little effect on average adult survival since *<* 0.5% of adults reached that age. Mate fidelity was set to zero, implying a single father per breeding season and no multiple pairings for males (in CKMRpop) and hence no within-cohort half-sibling pairs.

Samples were taken from the simulated populations and kinship computed using `CKMRpop` at sampling proportions 0.25%, 0.35%, and 0.45% representing small, medium and large samples of the population. Sample sizes were on average close to 290 for 0.25%, 410 for the 0.35%, and 540 for the 0.45% scenario. The average number of HSPs per sampling scenario was approximately 60, 130, and 230 respectively. Each individual in the sample was given a simulated noisy age score based on four linear and segmented regression models.

For the full-marginalisation model, we allowed abundance to vary over time via 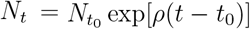, where *ρ* is the population rate of increase. This extends the constant *N* model of Section 2.2 and is included as an additional element of 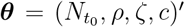. In the simulation, the population was stable (*ρ* ≈ 0).

The first scenario assumes a linear relationship with normal error, i.e., *G*_*i*_ = *α* + *βa*_*i*_ + _*i*_, where *α* = 0, *β* = 1 and _*i*_ ∼ *N* (0, *σ*^2^) with *σ* = (0.5, 1, 2), which simulated low to high variance in age score measurements given chronological age. This regression mimics the most reliable part of the relationship between chronological age and vertebral band count for school sharks — ages between 0 and the age at maturity (≈ 11) are considered linear with slope 1 and an intercept at 0 (Walker et al., 2001). The second simulation shows that the estimation method can scale arbitrarily so we repeat this linear simulation with *α* = 20, *β* = 10 and *σ* = (2, 4, 8).

The third simulation scenario uses a segmented regression model *G*_*i*_ = *α* + *β*_1_*a*_*i*_ + (*β*_2_ − *β*_1_)(*a*_*i*_ − *ψ*)_+_ + *ε*_*i*_ where *α* is the intercept, *β*_1_ is the slope of the line before the breakpoint, *β*_2_ is the slope after the breakpoint, *ψ* is the breakpoint, (*a*_*i*_ − *ψ*)_+_ = max(0, *a*_*i*_ − *ψ*) is the hinge function, and *ε*_*i*_ ∼ *N* (0, *σ*^2^) is the normal random error term. In the segmented regression analysis we set *β*_1_ = 1, *β*_2_ = 0.36 and *ψ* = 10. The motivation for this model was that it is hypothesised for school shark that the rate at which vertebral bands are deposited slows after maturity. A vertebral band deposit rate of 0.36 after age 11 was the value assumed for the first CKMR study of school shark (Thomson et al., 2020).

The fourth simulation investigated the potential of noisier measurements at higher ages due to varying individual rates of vertebral band deposition and increasing difficulty in band detection. We simulated a Gamma generalised linear model with *G*_*i*_|*A*_*i*_ ∼ Gamma(*µ*_*i*_, *φ*_disp_) where *µ*_*i*_ = *f* (*A*_*i*_) is the segmented regression formula (identity link) and *φ*_disp_ is the dispersion parameter (differentiated from survival) of the gamma distribution. The gamma distribution further models the positive measure of vertebral band count. We set *φ*_disp_ to (0.005, 0.01, 0.02), which corresponds to a variance of (0.5, 1, 2) at age 10 under the 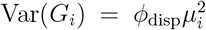 structure. We constrain the mean function to be positive during the fitting process by fitting *α* as the log parameter and *β*_1_ and *β*_2_ are bounded below at 0 but rarely near that value in fitting given their simulated values.

For each simulation scenario, 50 replicates were performed. For the full-marginalisation model we assumed a bin size *l* = 2 for the vertebral band age scores. The breakpoint was fixed at its true value in the segmented regression scenarios.

Initial pilot work indicated that in low sample size scenarios the model found it challenging to separate *σ*^2^ from the total variance, suggesting that age-score variance was difficult to reliably estimate from the CKMR data alone. We therefore constrained *σ* and *φ*_disp_ through a Gaussian prior on the log scale with expectation at the log of the true simulated value and standard deviation of log(1.1). To investigate sensitivity to this prior, we created a new scenario by altering the assumed mean in the scaled linear scenario, setting it to 3 and 6 for the *σ* = (2, 4) cases and to 4 for *σ* = 8.

### 3.2 Simulation results

In the linear model scenarios, the full-marginalisation model consistently outperformed the pairwise difference model. Both estimated *β* close to unity, but the pairwise difference model showed increasing upward bias as error variance grew, whereas the full-marginalisation model remained stable across all scenarios (Table **??**). The full-marginalisation model also produced less biased estimates of population abundance (Figures **??** and **??**). Other demographic parameters were well recovered: mortality was close to the simulated value of 0.22, though marginally upwardly biased, and *ρ* was estimated near zero as expected. Prediction variance showed only modest reductions with increasing sample size, suggesting it is largely driven by the inherent variability in the age score given chronological age rather than parameter uncertainty (Figure 1). The scaled linear scenario (*α* = 20, *β* = 10) confirmed that the method generalises to arbitrary scale and location (Table **??** and Figures **??** and **??**). Estimates of *σ* remained close to their prior values across all scenarios, consistent with the limited ability to estimate age-score variance from CKMR data alone. However, the sensitivity analysis showed that other parameter estimates were reasonably robust to misspecification of the prior mean for *σ* (Table **??** and Figure **??**). Estimates of *c* were near unity across all scenarios, consistent with the simulation design in which within-cohort HSPs were absent; with no such pairs in the data, the likelihood is flat with respect to *c* and the estimate reflects the initialisation rather than an identifiable signal.

**Figure 1.**
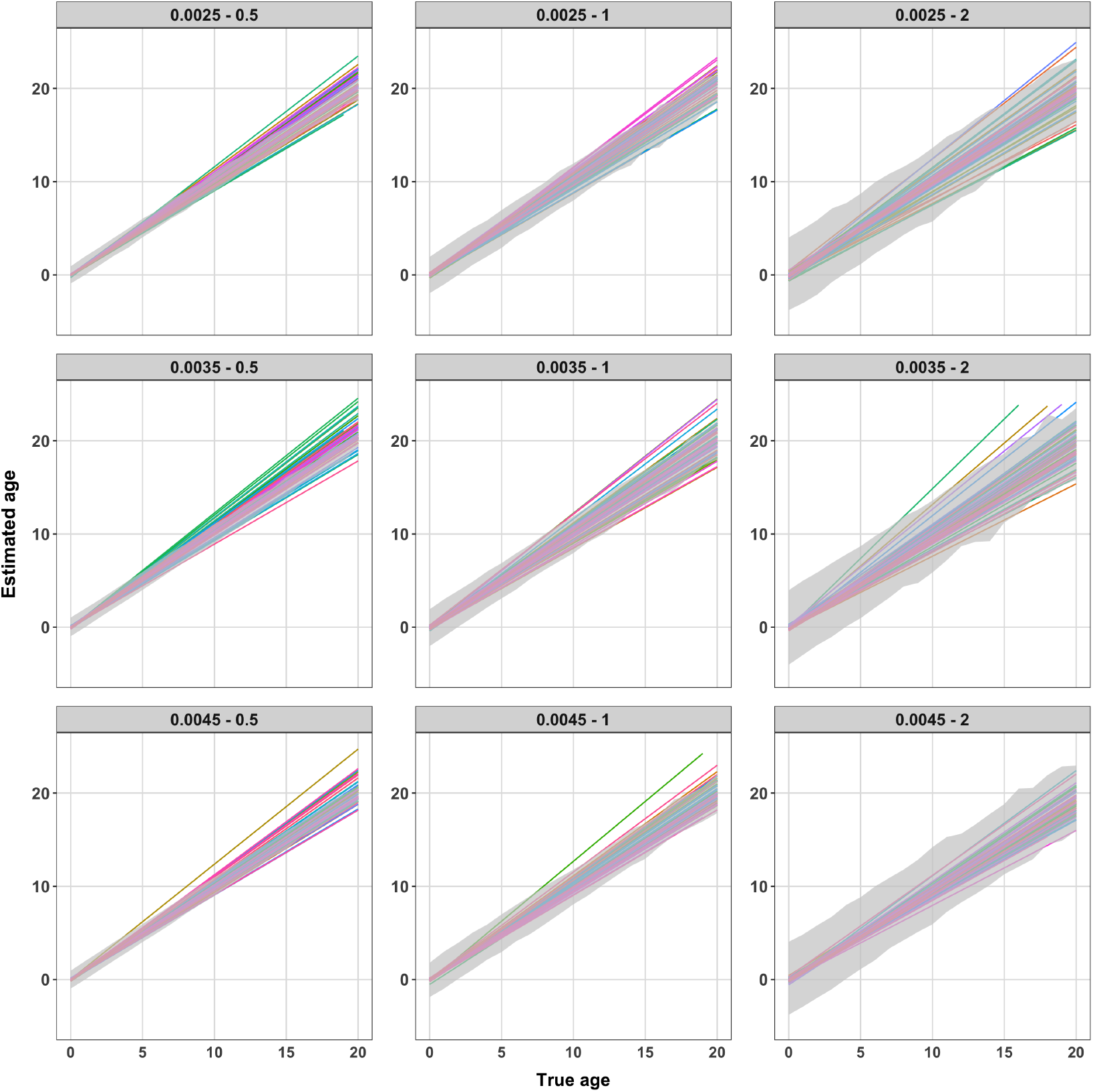
Summaries of estimated linear relationships from the full-marginalisation model in the linear model simulation scenario. Each panel shows increasing sample size from top to bottom and increasing error variance from left to right. Panel headings describe the sampling proportion and assumed standard error for the normal error term. The simulated relationship was *α* = 0 and *β* = 1. Each predicted line has a different colour and the grey transparent interval is the 95% percentile interval for the simulated noisy measure of age given the discrete chronological age value.

The segmented linear model scenarios tested the extension to non-linear relationships, including a gamma GLM with segmented linear predictors to allow for heteroscedastic errors. We report results only for the full-marginalisation model, as the pairwise difference approach does not extend naturally to non-linear relationships. For the segmented linear model with normal error, we found that estimates of the two slope parameters showed a small downward bias. We observed a partial correction of the bias with increasing sample size in high-variance combinations. High-variance scenarios also showed other demographic parameter estimates with bias, especially the mortality parameters (Table **??**). Population adult abundance estimates appeared largely well estimated, on average, in this scenario (Figure **??**).

Similarly, for the gamma GLM, we saw some downward bias in slope estimates but with less bias in high sample size combinations (Table **??**). Plots of the relationships between chronological and estimated age showed lines with mostly desirable properties including lower variance in estimates from high sample size and lower error variance combinations. Predicted linear relationships were nearly covered by the 95% percentiles computed from the simulation in each of the simulation combinations (Figure 2). Again, population adult abundance was well estimated (Figure **??**). Overall, the simulations showed that the full-marginalisation method recovers both the age score to chronological age relationship and key demographic parameters well across linear and non-linear scenarios, with performance degrading primarily in high-variance, low-sample-size combinations.

**Figure 2.**
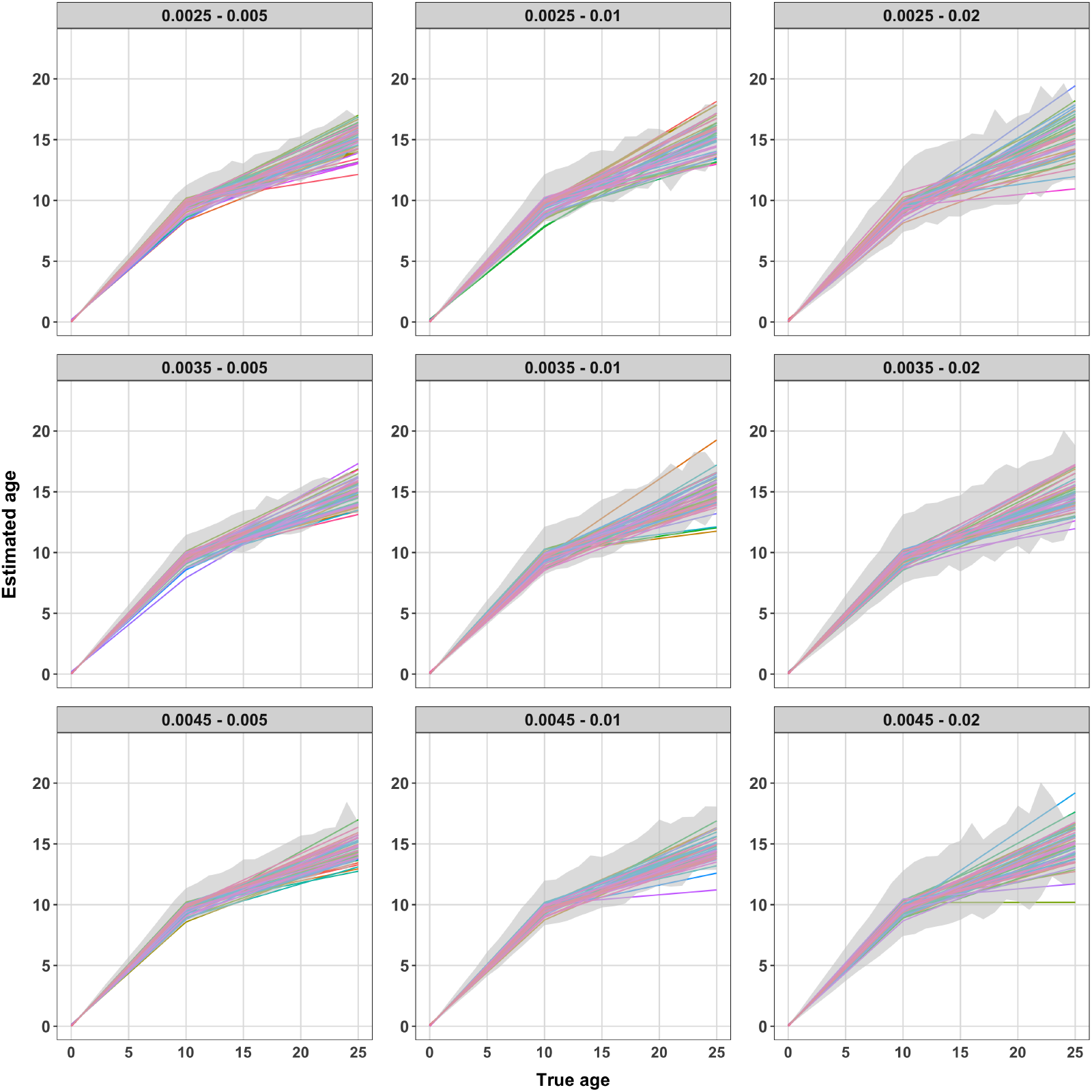
Summaries of estimated segmented linear relationships from the full-marginalisation model in the generalised linear model with gamma error. Each panel shows the results with increasing sample size from top to bottom and increasing error variance from left to right. Panel headings describe the sampling proportion and assumed standard error for the gamma error model. The simulated relationship was *α* = 0, *β*_1_ = 1, *β*_2_ = 0.36 and breakpoint at chronological age of 10. Each predicted line is from a replicate and has a different colour. The grey transparent intervals are the 95% percentile intervals from the simulated data, with the interval calculated for the set of noisy age values for a given discrete chronological age value.

## 4. Application to school shark vertebral band count

### 4.1 Data and model specification

The management of school shark in the Southern and Eastern Scalefish and Shark Fishery was historically based on a stock assessment model that relied on catch per unit effort (CPUE) as an index of abundance (Punt et al., 2000). However, due to changes in fishing practices and more restrictive management measures, CPUE became unreliable. As a result, alternative methods for monitoring the population were explored (Knuckey et al., 2014), leading to the adoption of close-kin mark-recapture (CKMR) with a first study completed in 2020 (Thomson et al., 2020). CKMR provides an estimate of absolute abundance using genetic data from related individuals rather than relying on CPUE, making it more robust to changes in fishing practices.

The data used in this application were acquired from a CKMR study for school shark that took samples from the fishery in southern Australia (Thomson et al., 2020). Detailed methods are available in the work of Thomson et al. (2020), but briefly a total of 3,035 school shark samples were collected from three sites in southern Australia. Quality control measures were applied, eliminating low-quality DNA samples, duplicate samples, and 13 misidentified gummy sharks. After processing, 2,424 high-quality samples remained for genetic analysis. Genotyping was performed for all samples using technology developed by Diversity Arrays Technology (DArT Pty Ltd, Canberra). The methods and results of genotyping and kin finding are further detailed in Thomson et al. (2020). Overall, the genotyping and kin finding processes worked well for school shark, and with little ambiguity regarding the identification of kin pairs. Of interest to auto-calibration are the 65 half-sibling pairs found in the study of Thomson et al. (2020). We used the data “as prepared” for that study to investigate the relationship between observed age and chronological age for school shark.

Ageing of school shark is based on counting hyper-mineralised bands in vertebrae. The Fish Ageing Service Pty Ltd (FAS, Queenscliff, Australia) aged the school shark samples, with a maximum of 26 bands counted (Figure **??**). While most samples were expected to be younger than 11, the fishery shifted towards longlines to avoid sea lions, resulting in a greater number of older sharks sampled. Walker et al. (2001) validated vertebral bands for aging using tagging studies and oxytetracycline, which binds to mineralising hard tissue, and provides a permanent mark in the vertebra for later comparison with time at liberty. This validation implied that vertebral bands correspond well up to 11 years but underestimate chronological age beyond that (Walker et al., 2001). Kalish (2002) used bomb radiocarbon dating to confirm underestimation of older ages but did not provide an alternative deposition rate.

For school shark, vertebral band counts were binned into discrete intervals (*l* = 2 years) to facilitate analysis. All pairwise comparisons were made and HSP relationships marked. Each shark pair was ordered by approximate birth year based on vertebral age score. Input data included the tabled combinations for each sampling year/vertebral band count bin combination and the associated number of kin within each bin. For the zero-year-old individuals (neonates) in the sample we assumed them to have a vertebral age score of zero — this information was included through a regression log-likelihood added to the CKMR auto-calibration log-likelihood. A segmented regression model was used with both the normal and gamma error structures. The permissible range of chronological ages was set to 0 to 35. The standard error and dispersion parameters were constrained on the log scale using a normal prior with mean value of log(0.9) for the normal model and log(0.05) for the gamma model — for both models a standard deviation of log(1.2) was used for the normal prior. These values were informed by a separate assessment of full-sibling pairs, which for school shark must be nearly always from the same birth cohort (34 FSPs were used with vertebral band counts up to 11 years (results not shown)). Population abundance was assumed stable for the analysis (*ρ* = 0). The segmented regression change point was fixed at *ψ* = 11, which is the approximate age at maturity. Fixed and random design matrices, and the thin-plate spline smoothing penalty matrix, for the GAM smooth estimation of the chronological-age priors were obtained using `mgcv`. Optimisation was performed with Template Model Builder (TMB) (Kristensen et al., 2016) and `nlminb` in R with reasonable lower bounds on parameters. The model outputs included model parameter estimates and uncertainty, the estimated smooth functions for the age distributions in each sampling year, adult abundance over sampling period, and the relationship between vertebral and chronological ages, with uncertainty quantified via standard errors and Monte Carlo simulations (10,000 draws). Visualised results included predicted versus observed age frequencies and the vertebral-to-chronological age relationship with confidence and prediction intervals.

### 4.2 Results

For the processed school shark data we ran the full-marginalisation auto-calibration for both the segmented linear model with normal error and the gamma GLM extension. Results from both models were similar across many of the main parameters of the model with adult abundance showing an interval estimate of 57,000 to 165,000 with central mass at 93,000 to 100,000 (Table 1 and Figure 3). Estimates for the relationship between chronological age and vertebral band count showed some difference between models with slopes marginally below unity for the normal model and above for the gamma GLM (Table 1). The normal model estimated the intercept to be near one, which is likely due to the positive nature of the vertebral band count and the symmetric error of the normal model. The intercept estimate from the gamma model was close to zero. Prediction intervals from both models showed the plausible spread of vertebral band counts for each chronological age value; both models showed that at chronological age 10/11 plausible vertebral band count values included those expected if the slope was truly one. The normal and gamma models estimated a slope substantially less than one after age at maturity (Figure 3). Smooth estimates of the chronological age distribution are similar between models and show plausible shapes. The estimated vertebral band count distributions for each sampling year matched the overall shape of observed distributions for both models but missed some within-year peaks in sampling for vertebral band counts between 5 and 10 years. Samples for years 2012 and 2014 were absent and low respectively, hence the flattened smooth estimates.

**Table 1.** Parameter estimates from full-marginalisation model applied to school shark. The intervals in brackets are 95% confidence intervals.

| Parameter | Normal estimate | Gamma estimate |
| --- | --- | --- |
| $N_{\text{adult}}$ | 100,800 (61,600, 165,200) | 93,800 (57,100, 154,200) |
| $\alpha$ | 1.09 (0.91, 1.26) | 1e-04 (1e-5, 1e-3) |
| $\beta_1$ | 0.94 (0.79, 0.95) | 1.14 (1.08, 1.20) |
| $\beta_2$ | 0.43 (0.40, 0.47) | 0.23 (0.14, 0.31) |
| $\zeta$ | 0.05 (0.02, 0.13) | 0.06 (0.025, 0.16) |
| $\sigma/\phi$ | 0.87 (0.74, 1.03) | 0.03 (0.02, 0.05) |
| $c$ | 0.65 (0.04, 11.4) | 0.60 (0.08, 4.40) |

**Figure 3.**
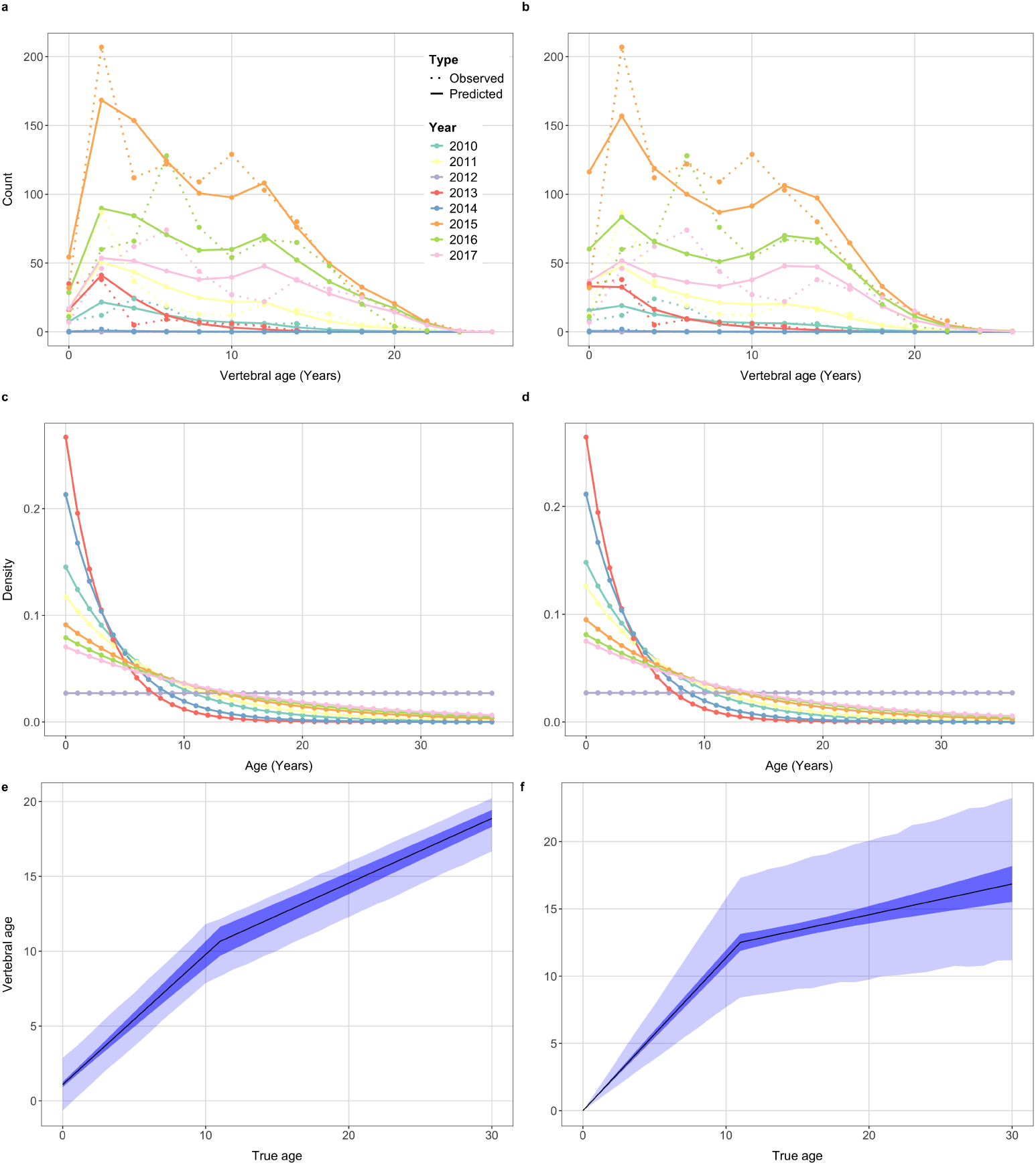
Model summaries for school shark for the segmented linear model with normal (left) and gamma (right) error. Panels a) and b) show the observed (dashed) and predicted (solid) distributions of vertebral band count for each sampling year. Panels c) and d) display the smooth prediction for the distribution of chronological age for each sampling year; a flat line is predicted for 2012 as there were no sampled school shark in that year. Panels e) and f) show the estimated relationship between true and vertebral band count with 95% confidence intervals and 95% prediction intervals derived from percentiles of Monte Carlo draws.

## 5. Discussion

We have shown that it is possible to estimate the relationship between age score and chronological age directly from multi-year CKMR HSP data without any need for known-age samples. Furthermore, this can be done without increasing the sample size beyond what is needed for CKMR itself. This novel method widens the scope of viable species for CKMR. Our ultimate goal is epigenetic age scores derived from methylation data, but the approach is also useful for calibrating uncertain age measurements such as vertebral or otolith band counts, and potentially even for estimating growth curves if the score is length. We presented a simple low-noise model for exposition, and extended this to an application-ready full-marginalisation model that accounts for age-score noise but requires a prior for chronological age. In simulation, the full-marginalisation method recovered both linear and non-linear age-score relationships, estimated variance and non-normal error structures, and produced reliable demographic parameter estimates. For school shark, the estimated relationship between chronological age and vertebral band count was aligned with expectations: approximately one-to-one until age at maturity, with evidence for a substantial slowing of the vertebral band deposition rate thereafter, supporting prior hypotheses. Demographic parameter estimates were concordant with the prior CKMR study of Thomson et al. (2020).

Errors-in-variables (here, age) are a well-known source of bias in demographic models, including CKMR (Petersma et al., 2024). Measurement error in age or size can lead to substantial overestimation of population growth rates through regression dilution (Louthan and Doak, 2018), and previous approaches to mitigating this have relied on auxiliary information from known-age individuals (Conn and Diefenbach, 2007). Our method overcomes this by coupling kinship information with capture times to estimate the age-score relationship directly, incorporating this uncertainty into demographic inference without requiring known-age samples.

The simulation results showed that the full-marginalisation model can overcome these biases and return reliable demographic parameter estimates.

There are several practical considerations in implementation. First, the age score must be discretised into bins, requiring a choice of bin width; in our experience, convergence was not sensitive to natural choices (e.g., 1- or 2-year bins for school shark). Second, the full-marginalisation model requires a prior distribution for chronological age in each sampling year. When the age score is precise, this prior has little practical influence because chronological ages are then tightly constrained by the observed score, but it remains a necessary component of the formulation. We presented a general solution in which this prior is estimated from the marginal distribution of age scores by discrete deconvolution within the overall model. Although this performed well in our examples, it is relatively elaborate, especially since the resulting priors were smooth and close to exponential distributions. In many applications, simpler alternatives may therefore be adequate, such as an exponential prior with rate estimated within the model, or a fixed prior assessed through sensitivity analyses. Where an age-structured population model already exists, the prior on chronological age could instead be derived directly from quantities already available in that framework.

A separate practical issue is estimation of the age-score variance, *σ*^2^. Supplementary Note **??** shows that *σ*^2^ is theoretically identifiable under a stable-age model with normal errors, but in practice the information available from CKMR data alone is often weak, particularly at moderate sample sizes. Biological structure can provide some additional information. For species that produce only one offspring per year, maternal half-sibling pairs with birth-gap 0 are impossible; more generally, if reproduction occurs only every *x* years, then certain birth gaps are excluded. School sharks are believed to reproduce non-annually, with a pupping interval of about three years. We did not model this cycle explicitly because our goal was age estimation from close-kin information rather than inference on annual reproductive state or demographic output. Nonetheless, the periodicity provides indirect information because larger age-score variance increases the frequency with which observed pairwise age-score differences are compatible with biologically impossible gaps. In many settings, however, auxiliary data will likely be more useful for estimating *σ*^2^. For school shark, we used full-sibling pairs known to share the same birth date, even though their absolute ages were unknown; analogous information could come from known-age individuals at selected ages, or from repeated age-score measurements on recaptured individuals separated by known time gaps. More formal joint models could also be developed, for example by combining an auto-calibration likelihood with a mark-recapture model for full-sibling pairs and shared age-score parameters. Overall, only a modest amount of effectively known-age information may be needed to estimate *σ*^2^, but some prior or auxiliary information will often remain important in practice.

The application of our auto-calibration method to school shark vertebral band count data demonstrated its practical utility. The method estimated a deposition rate consistent with one band per year until age at maturity, with a dramatic slowing post-maturity to between 0.14 and 0.47 bands per year across the two models fitted. These results corroborate previous findings: tag-recapture studies estimated that vertebral band counts correspond well with chronological age up to approximately 11 years but systematically underestimate chronological age beyond that point (Moulton et al., 1992), with Walker et al. (2001) estimating a post-maturity deposition rate of 0.36 bands per year. Bomb radiocarbon dating confirmed the underestimation of older ages but provided no estimate of the deposition rate (Kalish, 2002). The gamma model provided a more plausible fit than the normal model, as expected given the strictly positive data, with an intercept estimate close to zero. Overall, the estimates validate the method’s ability to recover known age-score relationships and integrate age uncertainty into a CKMR model.

Looking ahead, the most promising application of auto-calibration may be in combination with minimally invasive sampling techniques. A single tissue biopsy can yield both genotype data for kinship inference and DNA methylation data as an age score (Peters et al., 2023), enabling joint inference of age and demography without known-age validation samples. This is particularly relevant given the difficulty of obtaining reliable chronological age data for wild populations, which continues to limit the application of epigenetic clocks (Hanski et al., 2024). Extending the framework to handle more complex error structures, non-linear age-score relationships beyond segmented regression, and richer population dynamics models would further broaden its applicability across taxa.

## Acknowledgements

The authors would like to sincerely acknowledge Floriaan Delva-Devloo and Rasanthi Gunasekera for their help with sample processing and logistics. We thank the Fisheries Research and Development Corporation for funding the CKMR project that collected the school shark samples and Simon Robertson from Fish Ageing Services Pty Ltd for their great help with the vertebral band ageing.

## Data and code availability

Simulation and school shark application code is available at https://github.com/lukelloydjones/Validation-free-age-calibration. School shark data underlying the analysis will be deposited to the permanent Data Access Portal (DAP, https://data.csiro.au/) repository on acceptance.

## 1 Supplementary Notes

### 1.1 Identifiability of parameters of pairwise difference model

Let 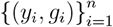 be observed pairs of discretised year and age scores, and for each pair 1 ≤ *i < j* ≤ *n*, define Δ*y*_*ij*_ = *y*_*j*_ − *y*_*i*_ and Δ*g*_*ij*_ = *g*_*j*_ − *g*_*i*_, which are realisations of *δy*_*ij*_ and *δg*_*ij*_. Let *x*_*ij*_ ∈ {0, 1} be the observed indicator that the pair is MHSP. Assume the pairwise-difference model (described by (5) in the main text) holds for all *i* and *j*:

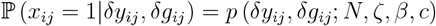

where

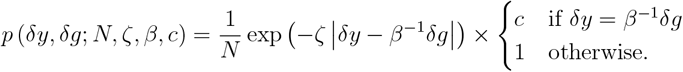

Fix contrasts *t*_0_ = (0, 0), *t*_1_ = (*d*_1_, 0), *t*_2_ = (*d*_2_, 0), and *t*_3_ = (0, *g*_0_), where *d*_1_, *d*_2_ ∈ N are distinct, and *g*_0_ = 0 is a nonzero score difference. For each *t* = (*δy, δg*), define the set of pairs matching that contrast and its size by

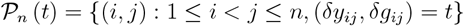

and *m*_*n*_ (*t*) = |𝒫_*n*_ (*t*)|. Define the number of observed MHSPs in that cell by 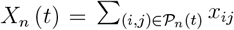 and write the empirical conditional MHSP rate (when *m*_*n*_ (*t*) *>* 0) by

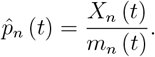

Suppose that there exists a true set of parameters (*N*_0_, *ζ*_0_, *β*_0_, *c*_0_) such that the following assumptions hold:

**A1** For each *r* ∈ {0, 1, 2, 3}, *m*_*n*_ (*t*_*r*_) → ∞ in probability (i.e., you can sample infinitely many observations of each contrast)

**A2** Conditionally on 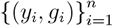, the indicators {*x*_*ij*_ : (*i, j*) ∈ 𝒫_*n*_ (*t*_*r*_)} are independent Bernoulli random variables with mean *p* (*t*_*r*_; *N*_0_, *ζ*_0_, *β*_0_, *c*_0_).

#### Proposition 1

*Under A1 and A2 and the setup above:*

1. *For each r* ∈ {0, 1, 2, 3}, 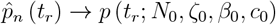 *in probability*.
2. *The map:*

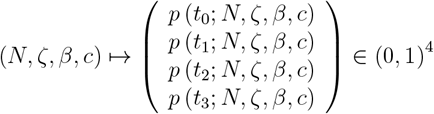

*is one-to-one, hence the four contrast point conditional probabilities uniquely determine* (*N, ζ, β, c*).

*Proof*. (1) Fix a contrast *t* = *t*_*r*_. Conditional on the observed {(*y*_*i*_, *g*_*i*_)}, the random variables {*x*_*ij*_ : (*i, j*) ∈ 𝒫_*n*_ (*t*)} are i.i.d. Bernoulli with success probability *p* (*t*; *N*_0_, *ζ*_0_, *β*_0_, *c*_0_), thus

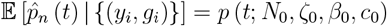

and

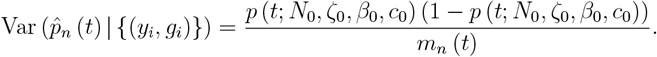

By Chebyshev’s inequality, for every *E >* 0,

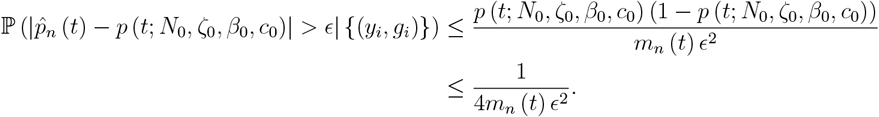

Using (A1) then yields the desired result.

(2) Assume that there are two parameter vectors (*N, ζ, β, c*) and (*N* ^t^, *ζ*^t^, *β*^t^, *c*^t^) such that

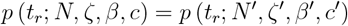

for each *r* ∈ {0, 1, 2, 3}. Using the explicit form of *p*, for *t*_1_ = (*d*_1_, 0) and *t*_2_ = (*d*_2_, 0), *d*_1_ /= *d*_2_, the lucky-litter factor is inactive so we have:

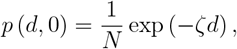

for *d* ∈ {*d*_1_, *d*_2_}. Hence,

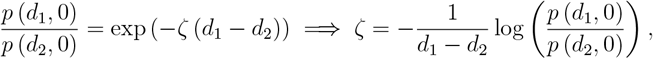

so *ζ* = *ζ*^t^. With *ζ* = *ζ*^t^, from *p* (*d*_1_, 0) = *N* ^−1^ exp (−*ζd*_1_), we have

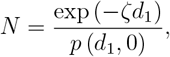

Next, from *t*_3_ = (0, *g*_0_) with *g*_0_ = 0, the lucky-litter factor is again inactive and we have

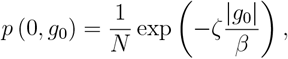

but since *N* = *N* ^t^ and *ζ* = *ζ*^t^, this forces *β* = *β*^t^. Finally, with *t*_0_ = (0, 0), the lucky-litter factor is in effect and we have *p* (0, 0) = *N* ^−1^*c*, thus because *N* = *N* ^t^, we must have *c* = *c*^t^, yielding the desired result.

### 1.2 Formalisation of collapse of full-marginalisation to pairwise difference approximation

#### Proposition 2

*For fixed index pair* (*i, j*), *write:*

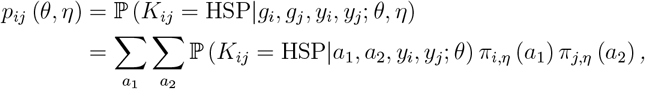

*where π*_*i,η*_ (*a*) = ℙ (*A*_*i*_ = *a*|*g*_*i*_, *y*_*i*_, *s*; *η*) *and similarly for π*_*j,η*_. *Assume that there exists a sequence of age-measurement parameter vectors* (*η*_*m*_)_*m*_ *and deterministic* 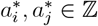, *such that*

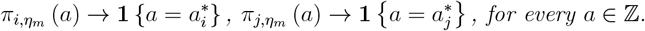

*Then*,

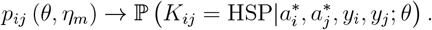

*Moreover, under the special case of (3) with E*_*i*_ = 0,

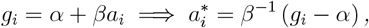

*and under the special case of (2) with b* = *y* − *a, the limit recovers the pairwise difference form described by (5) in the main text:*

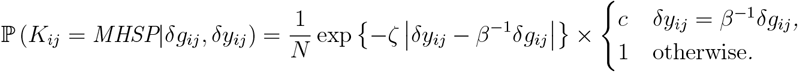

*Proof*. Let

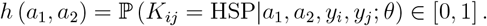

Then,

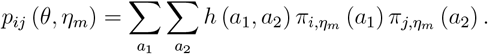

Write

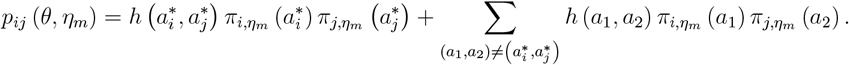

Since 0 ≤ *h* ≤ 1, it follows that

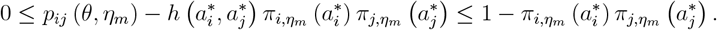

Since 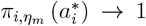 and 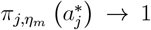, we have 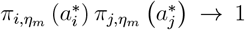, and thus 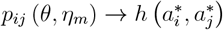, as required.

### 1.3 Discrete deconvolution via smoothing

Equation (6) of the main text requires the double sum over ages for each pair of observed (*g*_1_, *g*_2_) values. Because the values of *g*_*i*_ are continuous this would require a unique sum for all pairs. To improve computational efficiency we discretise the range [*c, d*] of observed *g* values into bins *q* ∈ (1, …, *Q*) of interval size *l* chosen by the modeller and compute the probabilities for the midpoint values of each grid cell. We check whether the resolution is too coarse by altering *l* until the parameter estimates change by some small tolerance. This computation is preferable as it is quadratic in the number of grid cells rather than the sample size.

For bins of *g*, we can define a matrix **M** with entries *M*_*ag*_ = ℙ (*G* = *g a*; ***η***) so that each row of **M** corresponds to the distribution of observed age scores given chronological age. The distribution of the ages of sampled individuals in year *y* is denoted ℙ (*A* = *a*|*y, s*) = *f*_*y*_(*a*) and modelled as a smooth function of age. Therefore, for discrete ages and binned scores the problem is to estimate some smooth function for each sample year such that,

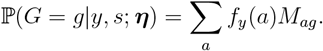

Let ***C***_*y*_ be a vector of *g* values over all *q* bins for a given sampling year, 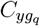 is equal to the number of *g* values in the sample counted in bin *q*. Then the random variable *G* for a given *y* is given by *G* ∈ {*g*_1_, …, *g*_*Q*_} in which the probability of a realised discrete *g* value is denoted 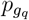. Suppose we observe a total of *n*_*y*_ values of *g* in year *y*, it follows that the count vector for a given sampling year can be modelled with a multinomial distribution,

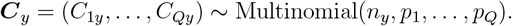

Alternatively, this can be modelled as independent Poisson distributions conditional on the sum over all bins being Poisson, by noting a useful relation between the multinomial and Poisson distributions (Steel, 1953). We provide a proof to complete the connection in Section 1.4.

We prefer the maximisation of a Poisson log-likelihood because it is computationally easier and more stable than for a multinomial distribution.

Based on the above discussion we elect to model the counts for each discrete *g* score as independent Poisson random variables

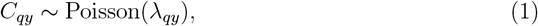

where the total sum of all counts of *g* in *Q* bins in year *y, N*_*y*_ = Σ_*q*_ *C*_*qy*_, is now a random variable. We parameterise the Poisson rates as *λ*_*qy*_ = *µ*_*y*_*p*_*qy*_, where *µ*_*y*_ denotes the expected total number of observations in year *y*, and

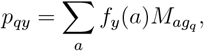

is the modelled probability of observed score *g*_*q*_. Using the fact that the expected value of a Poisson random variable is simply its rate parameter we have,

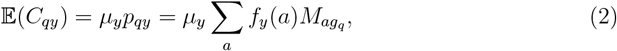

where *f*_*y*_(*a*) is the smooth function defined above. To estimate the smooth function we use the principles of generalised additive models. Computationally, each sampling year is assigned a set of random effects *u*_*yk*_ such that,

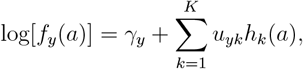

where *γ*_*y*_ is the intercept parameter. We choose *K* basis functions which are treated as known and have some unknown parameters **u**_*y*_ = (*u*_*y*1_, …, *u*_*yK*_)^t^. To control smoothness, we penalise “wiggliness” by choosing a basis dimension fixed at a size larger than believed necessary and introduce a penalty to the fitting process. We incorporate the belief that the true function is more likely smooth than wiggly by defining a prior distribution on function wiggliness such that

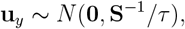

where **S**^−1^ is defined as the pseudo-inverse of **S**, and **S** is a positive semi-definite matrix depending on the basis functions (Wood, 2017). The parameter *τ* controls the strength of regularisation on the spline coefficients, with larger values inducing smoother estimates of *f*_*y*_(*a*).

We then estimate the posterior mean/mode **û**_*y*_ that maximises

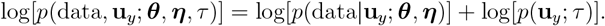

This structure allows for estimation via marginal likelihood maximisation with the integration over the random effects performed by Laplace approximation in TMB. The first component of the model includes the contributions from the CKMR data generating model that is primarily concerned with estimating ***θ***, which we treat as binomial for each (*y, g*_*q*_) combination with success probability defined by Equation (6) of the main text. Part 1 also includes the process for generating the distribution for binned *g* values in each sampling year, which are modelled as Poisson with rate parameters defined in Equation (2).

The elements required for the smoothing, which include the basis function matrix **H** where *H*_*ik*_ = *h*_*k*_(*a*_*i*_) and **S**, are generated by the mgcv package (Wood, 2011) in the R programming language. The default thin plate regression splines are used. Optimisation of the pseudo-likelihood estimates of ***θ, η*** and *τ* from both models is performed using the Template Model Builder (TMB) (Kristensen et al., 2016) package and nonlinear quasi-Newton BFGS optimisation routines implemented in the nlminb function from the stats package in R. TMB calculates first- and second-order derivatives of the likelihood function by algorithmic differentiation, and asymptotic standard deviations of any parameter, or derived parameter, are obtained by the delta method.

### 1.4 Proof of Poisson connection to multinomial

The following result is well known (Steel, 1953) but we provide a proof here for completeness with our notation and easy reference.

#### Proposition 3

*Fix a sampling year y. Let*

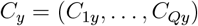

*and* 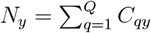. *Further let p*_*qy*_ ≥ 0 *satisfy* 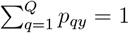. *Then:*

1. *If N*_*y*_ ∼ Poisson (*µ*_*y*_) *and C*_*y*_|*N*_*y*_ ∼ Multinomial (*N*_*y*_; *p*_1*y*_, …, *p*_*Qy*_), *then C*_1*y*_, …, *C*_*Qy*_ *are independent and for each q* ∈ {1, …, *Q*},

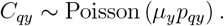
2. *If C*_*qy*_ *are independent with C*_*qy*_ ∼ Poisson (*λ*_*qy*_) *and* 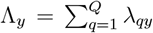, *then N*_*y*_ ∼ Poisson (Λ_*y*_) *and*

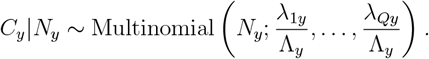
3. *With* 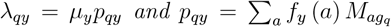, *for fixed N*_*y*_ = *n*_*y*_, *maximising the independent Poisson likelihood in p*_*qy*_ *is equivalent to maximising the multinomial likelihood, conditional on N*_*y*_ = *n*_*y*_.

1. *Proof*. (1) Fix nonnegative integers *c*_1_, … *c*_*Q*_ and let 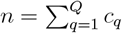. Then the product rule implies

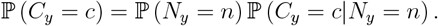

Substitute in the Poisson PMF and multinomial PMF:

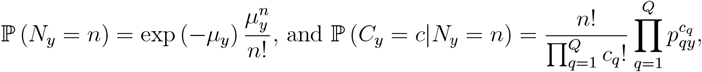

we get

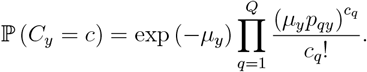

Since 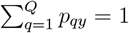, we can rewrite 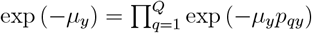, giving us

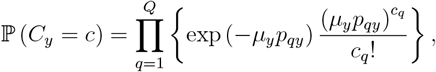

which is in the form of a product of Poisson PMFs. Thus *C*_*qy*_ are independent Poisson random variables with means *µ*_*y*_*p*_*qy*_.
2. If *C*_*qy*_ are independent Poisson (*λ*_*qy*_), then

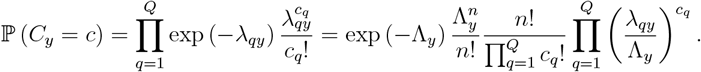

The first factor is P (*N*_*y*_ = *n*) for *N*_*y*_ ∼ Poisson (Λ_*y*_), and the second factor is the multinomial PMF. Dividing through by P (*N*_*y*_ = *n*) yields the desired multinomial distribution.
3. Under the Poisson model λ_*qy*_ = *μ*_*y*_*p*_*qy*_, the log-likelihood is:

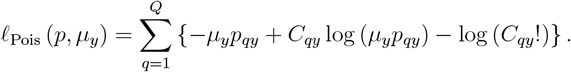

For fixed observed 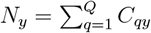, this becomes.

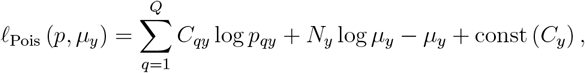

thus conditional on *N*_*y*_, maximising over *p* is equivalent to maximising Σ_*q*_ *C*_*qy*_ log *p*_*qy*_, i.e., the multinomial log-likelihood conditional on *N*_*y*_.

### 1.5 Non-identifiability of *α* in the marginalised likelihood

Let us fix measurement midpoints {*g*_1_, …, *g*_*Q*_} and bins *B*_*q*_ ⊂ R

(e.g., *B*_*q*_ = {*g* : *g*_*q*_ − */*2 *< g* ≤ *g*_*q*_ + */*2}), and define the discretised measurement as *G*^disc^ = *g*_*q*_ if *G* ∈ *B*_*q*_. Assume that (3) holds:

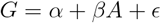

for *β >* 0, where *E* has density 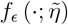 and write 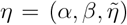. Then, as per the Supplementary Material 1.3, we discretise the measurement probabilities conditional on *A* as:

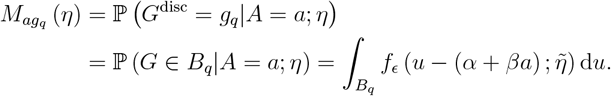

Next, for each sampling year *y*, let *f*_*y*_ (*a*) = ℙ (*A* = *a y, s*) be an unknown age PMF on Z, and define the implied discretised score distribution (i.e., the discretised version of (7)) as:

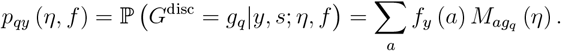

Further define the posterior on ages given discretised measurements (e.g., Bayes’ rule formula following (6)) by

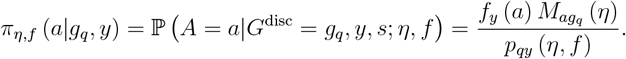

Lastly, assume that the HSP probability given age depends on (*a*_1_, *a*_2_, *y*_1_, *y*_2_) only through the birth-gap:

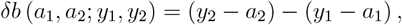

i.e., there exists a function *H*_*θ*_ : Z → [0, 1] such that

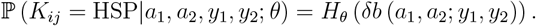

Then, the marginalised kinship probability (e.g., (6) with discretised *g*) is:

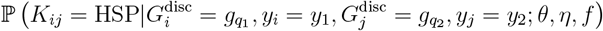

can be written as

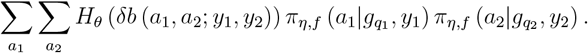

#### Proposition 4

*Under the setup above, for any fixed integer* Δ, *define the transformations* 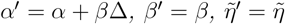, *and* 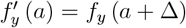, *for each y and a. Then, for every year y and bin q*,

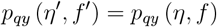

*and the posteriors satisfy the shift relationship:*

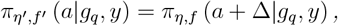

*and for every two cells* 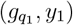 *and* 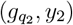,

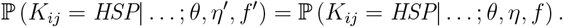

*Proof*. Firstly, observe that by substituting *α*^t^ = *α* + *β*Δ, we have

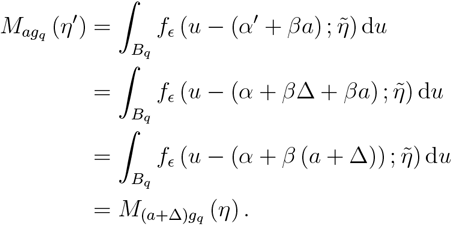

Then, it follows that

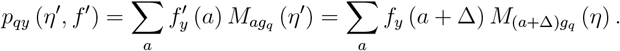

Making the change of variable *a*^t^ = *a* + Δ, we get

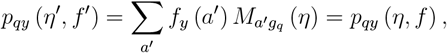

which proves the first statement. From Bayes’ rule, we then have

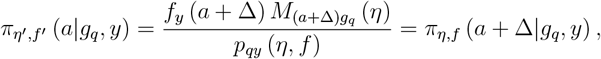

which yields the second statement. Lastly, starting from the discretised form of (6), substituting the equation above yields:

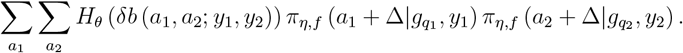

Then, making the change of variable 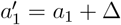 and 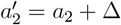, we have

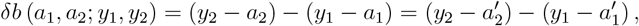

thus *H*_*θ*_ (*δb*) is unchanged, and the double sum is exactly the same as that under *η* and *f*, thus proving the third statement.

This implies that the likelihood induced by *η* and {*f*_*y*_} is invariant under shift and so *α* and the family {*f*_*y*_} are not identifiable without further constraint. This necessitates the additional likelihood elements corresponding to the known age individuals in order to estimate *α* and thus {*f*_*y*_}.

### 1.6 Identifiability of the parameters in the marginalised likelihood

#### Proposition 5

*Under the constant adult mortality assumed in our CKMR model, the stationary age distribution of adults is geometric. Thus, the distribution of chronological age A takes the form*

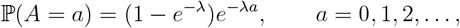

*with λ >* 0, *shared across all sampling years. Assume further that*

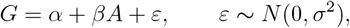

*where α is known, β >* 0, *and σ*^2^ *>* 0.

*For a sampled pair, let Y*_1_ *and Y*_2_ *denote the observable sampling years and, after relabelling the pair if necessary, define the observable sampling-year gap by*

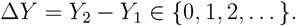

*Assume that the HSP probability depends on ages and sampling years only through the birth-gap*

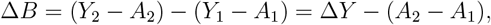

*via*

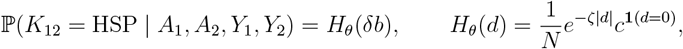

*with N >* 0, *ζ >* 0, *c >* 0. *Note that λ and ζ may differ if sampling is selective with respect to age; under random sampling from a stationary population, λ* = *ζ and the model simplifies accordingly*.

*Finally, assume that the support of the observable gap* Δ*Y contains* 0 *and at least two distinct positive values: that is, assume there exist integers* 0 *< d*_1_ *< d*_2_ *such that*

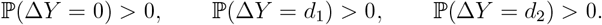

*Equivalently, the observable design includes same-year pairs (Y*_1_ = *Y*_2_*) and also pairs sampled at two different positive year gaps*.

*Then the parameter vector*

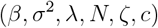

*is identifiable from the joint law of the pair-level observables*

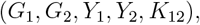

*or equivalently from the joint law of* (*G*_1_, *G*_2_, Δ*Y, K*_12_).

*Proof*. Write

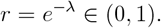

We first identify (*λ, β, σ*^2^) from the marginal score distribution, and then identify (*N, ζ, c*) from the kinship part. The support condition on Δ*Y* is used only in Step 3.

#### Step 1: identification of (*λ, β, σ*^2^) **from the score distribution**

The joint law of (*G*_1_, *G*_2_, *Y*_1_, *Y*_2_, *K*_12_) determines the marginal law of a single score, so it is enough to work with a generic score

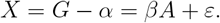

Because *A* ∼ Geom_0_(1 − *r*) on {0, 1, 2, … },

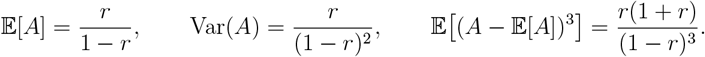

Since *ε* has mean 0, variance *σ*^2^, and third central moment 0,

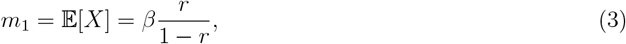

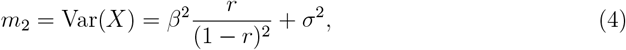

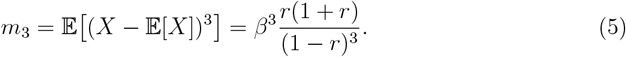

Therefore

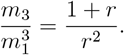

The right-hand side depends only on *r*, so *r* is identified as the unique positive solution of

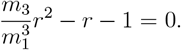

Hence *λ* = − log *r* is identified.

With *r* known, the first equation gives

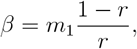

and then the second gives

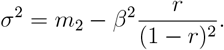

Thus (*λ, β, σ*^2^) are identified from the marginal distribution of *G*.

#### Step 2: very small scores correspond almost surely to age 0

For an observed score *g*, let

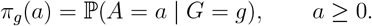

By Bayes’ rule,

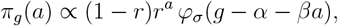

where *ϕ*_*σ*_ is the *N* (0, *σ*^2^) density. Hence, for each *a* ≥ 1,

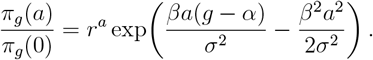

As *g* → −∞, the term *βa*(*g* − *α*)*/σ*^2^ tends to −∞, so

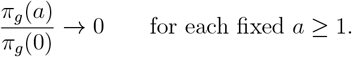

To conclude that *π*_*g*_(0) → 1, it is not sufficient to show that each ratio 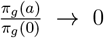 individually, since the sum of infinitely many vanishing terms may not itself vanish. We require that the sum Σ _*a*≥1_ *π*_*g*_(*a*)*/π*_*g*_(0) → 0, which by dominated convergence follows if each term is bounded above by a *g*-independent quantity that is summable over *a*. So, for all *g* ≤ *α*,

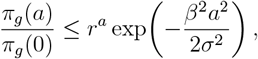

and the right-hand side is summable over *a* ≥ 1. Therefore the total posterior mass on ages *a* ≥ 1 tends to 0, and hence

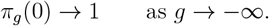

So conditioning on an extremely small score forces the age to be 0 with probability tending to 1.

#### Step 3: the support condition on Δ*Y* **lets us recover** *H*_*θ*_(0) **and** *H*_*θ*_(*δy*) **at two positive gaps**

Let *δy* be any value in the support of Δ*Y*, and define

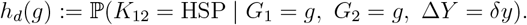

Because Δ*Y* = *δy* means that the two observable sampling years differ by exactly *δy*, the birth-gap becomes

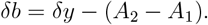

Hence, by conditioning on the two ages,

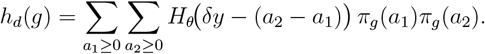

By Step 2, both posterior age distributions concentrate at 0 as *g* → −∞, so

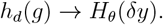

Thus, whenever *δy* is in the support of the observable gap Δ*Y*, the observable law determines the value *H*_*θ*_(*δy*).

Now use the explicit support assumption. Since ℙ (Δ*Y* = 0) *>* 0, same-year pairs are observed and therefore

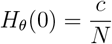

is identified.

Since ℙ (Δ*Y* = *δy*_1_) *>* 0 and P(Δ*Y* = *δy*_2_) *>* 0 for two distinct positive values *δy*_1_ /= *δy*_2_, the two positive-gap quantities

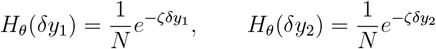

are also identified. Taking the ratio gives

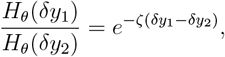

so

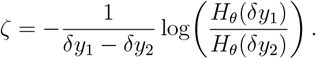

With *ζ* known,

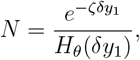

and then

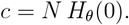

This also explains why the support condition on Δ*Y* is needed: same-year pairs identify *c/N* ; one positive gap would only identify the single combination *N* ^−1^*e*^−*ζδy*^; two distinct positive gaps separate *ζ* from *N* .

Therefore (*N, ζ, c*) are identified. Together with Step 1, this proves identifiability of

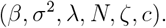

## Supplementary Tables

**Table S1.** Parameter estimates and standard deviations in brackets for pairwise difference and full-marginalisation models across nine simulation scenarios generated from the normal linear model. True simulated values are *α* = 0, *β* = 1, *ρ* = 0 and *σ* values are presented in each row. Estimates of *c* are expected to be near unity (see main text). A value 0.22 is expected for adult mortality from the simulated survival probabilities. The population was simulated to be on average stable i.e., *ρ* ≈ 0.

| Pairwise difference |  |  |  |  |  |  |  |
| --- | --- | --- | --- | --- | --- | --- | --- |
| Sample fraction | Model S.D. | | $\beta$ | $\zeta$ | | | $c$ |
| 0.0025 | 0.5 |  | 0.937 (0.335) | 0.216 (0.084) |  |  | 0.972 (0.339) |
| 0.0025 | 1.0 |  | 0.998 (0.397) | 0.199 (0.095) |  |  | 0.979 (0.430) |
| 0.0025 | 2.0 |  | 1.315 (0.526) | 0.189 (0.069) |  |  | 1.089 (0.426) |
| 0.0035 | 0.5 |  | 1.078 (0.214) | 0.235 (0.057) |  |  | 0.818 (0.256) |
| 0.0035 | 1.0 |  | 1.111 (0.278) | 0.220 (0.063) |  |  | 0.894 (0.357) |
| 0.0035 | 2.0 |  | 1.287 (0.418) | 0.178 (0.055) |  |  | 1.114 (0.337) |
| 0.0045 | 0.5 |  | 0.994 (0.259) | 0.228 (0.061) |  |  | 0.910 (0.312) |
| 0.0045 | 1.0 |  | 1.095 (0.287) | 0.212 (0.055) |  |  | 0.924 (0.258) |
| 0.0045 | 2.0 |  | 1.301 (0.258) | 0.173 (0.040) |  |  | 1.046 (0.176) |
| Full-marginalisation |  |  |  |  |  |  |  |
| | | $\alpha$ | $\beta$ | $\zeta$ | $\rho$ | $\sigma$ | $c$ |
| 0.0025 | 0.5 | -0.040 (0.078) | 1.029 (0.063) | 0.259 (0.082) | -0.003 (0.047) | 0.498 (0.024) | 1.427 (0.906) |
| 0.0025 | 1.0 | -0.045 (0.139) | 1.034 (0.068) | 0.248 (0.094) | -0.003 (0.046) | 1.007 (0.046) | 1.982 (1.488) |
| 0.0025 | 2.0 | -0.126 (0.271) | 0.998 (0.100) | 0.272 (0.146) | 0.007 (0.043) | 2.020 (0.100) | 3.746 (2.669) |
| 0.0035 | 0.5 | -0.057 (0.086) | 1.054 (0.075) | 0.254 (0.052) | -0.011 (0.032) | 0.501 (0.025) | 1.186 (0.503) |
| 0.0035 | 1.0 | -0.041 (0.144) | 1.022 (0.084) | 0.251 (0.060) | -0.002 (0.028) | 1.016 (0.049) | 1.758 (0.770) |
| 0.0035 | 2.0 | -0.071 (0.183) | 1.000 (0.127) | 0.259 (0.091) | -0.005 (0.032) | 2.043 (0.108) | 3.333 (2.231) |
| 0.0045 | 0.5 | -0.043 (0.073) | 1.013 (0.066) | 0.261 (0.038) | 0.003 (0.025) | 0.508 (0.030) | 1.217 (0.418) |
| 0.0045 | 1.0 | -0.036 (0.119) | 1.013 (0.067) | 0.243 (0.043) | -0.001 (0.029) | 1.032 (0.052) | 1.803 (0.680) |
| 0.0045 | 2.0 | -0.087 (0.203) | 0.959 (0.067) | 0.220 (0.064) | 0.006 (0.026) | 2.070 (0.111) | 3.725 (1.980) |

**Table S2.**
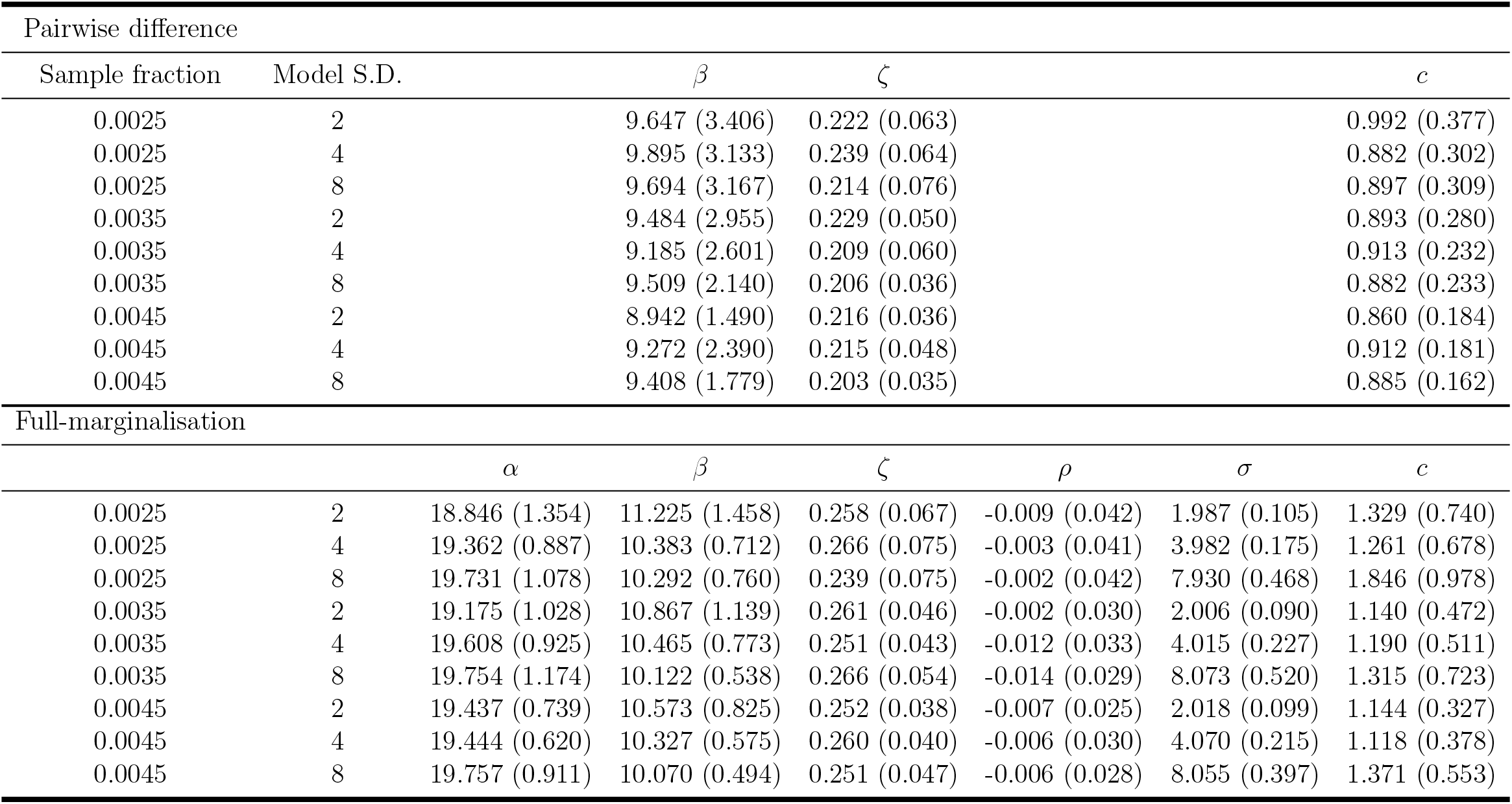
Parameter estimates and standard deviations in brackets for pairwise difference and full-marginalisation models across nine simulation scenarios generated from the scaled normal linear model. Simulated true values for *α* and *β* were 20 and 10 respectively. Estimates of *c* are expected to be near unity (see main text). A value 0.22 is expected for adult mortality from the simulated survival probabilities. The population was simulated to be on average stable i.e., *ρ* ≈ 0.

**Table S3.**
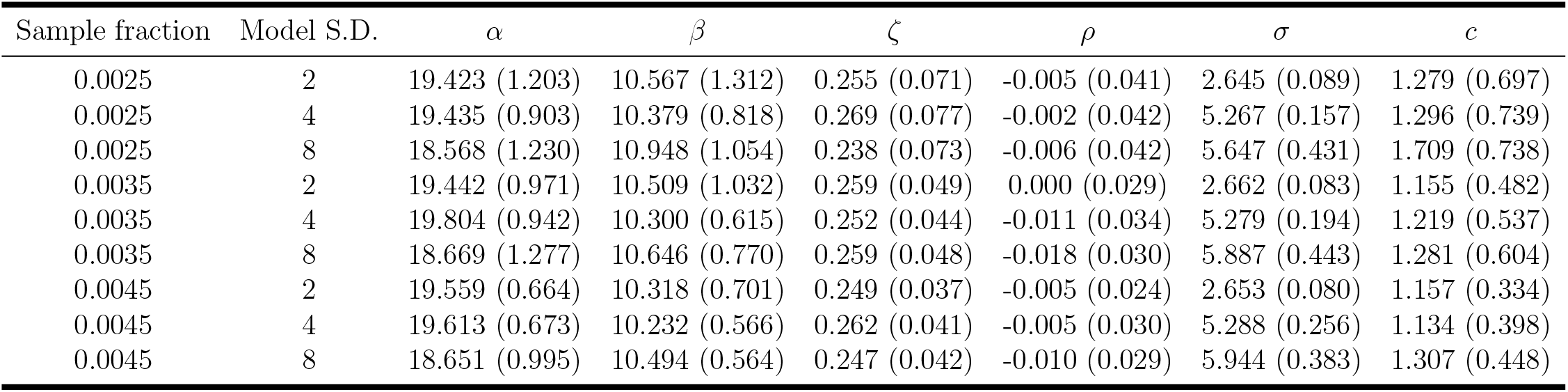
Parameter estimates and standard deviations in brackets for full-marginalisation with *σ* prior mean fixed away from simulated truth. For simulated model S.D.s of 2 and 4 the prior mean is set to 50% more and for 8 it is set to half that value. Results are from nine simulation scenarios generated from the scaled normal linear model. Simulated true values for *α* and *β* were 20 and 10 respectively. Estimates of *c* are expected to be near unity (see main text). A value 0.22 is expected for adult mortality from the simulated survival probabilities. The population was simulated to be on average stable i.e., *ρ* ≈ 0

**Table S4.** Parameter estimates and standard deviations in brackets for full-marginalisation models across nine simulation scenarios generated from the segmented linear normal error model. Simulated true values for *α, β*_1_ and *β*_2_ were 0, 1 and 0.36 respectively. Estimates of *c* are expected to be near unity (see main text). A value 0.22 is expected for adult mortality from the simulated survival probabilities. The population was simulated to be on average stable i.e., *ρ* ≈ 0.

| Sample fraction | Model S.D. | $\alpha$ | $\beta_1$ | $\beta_2$ | $\zeta$ | $\rho$ | $\sigma$ | $c$ |
| --- | --- | --- | --- | --- | --- | --- | --- | --- |
| 0.0025 | 0.5 | 0.012 (0.091) | 0.955 (0.059) | 0.330 (0.134) | 0.246 (0.057) | -0.005 (0.040) | 0.500 (0.024) | 1.225 (0.706) |
| 0.0025 | 1.0 | 0.005 (0.163) | 0.940 (0.106) | 0.354 (0.148) | 0.260 (0.082) | -0.012 (0.056) | 1.015 (0.053) | 1.879 (0.900) |
| 0.0025 | 2.0 | 0.336 (0.676) | 0.530 (0.435) | 0.543 (0.274) | 1.448 (4.515) | -0.004 (0.039) | 2.103 (0.123) | 2.370 (0.972) |
| 0.0035 | 0.5 | -0.026 (0.063) | 0.966 (0.039) | 0.341 (0.060) | 0.239 (0.051) | 0.001 (0.025) | 0.498 (0.027) | 1.354 (0.492) |
| 0.0035 | 1.0 | -0.008 (0.172) | 0.921 (0.084) | 0.359 (0.084) | 0.244 (0.062) | -0.002 (0.035) | 1.009 (0.066) | 1.932 (0.776) |
| 0.0035 | 2.0 | 0.113 (0.452) | 0.716 (0.332) | 0.450 (0.217) | 0.686 (2.802) | 0.001 (0.031) | 2.096 (0.145) | 2.510 (0.812) |
| 0.0045 | 0.5 | -0.025 (0.077) | 0.970 (0.037) | 0.325 (0.061) | 0.232 (0.036) | 0.001 (0.024) | 0.503 (0.027) | 1.360 (0.455) |
| 0.0045 | 1.0 | -0.032 (0.107) | 0.938 (0.071) | 0.331 (0.105) | 0.243 (0.054) | 0.005 (0.026) | 1.029 (0.054) | 1.735 (0.532) |
| 0.0045 | 2.0 | 0.092 (0.395) | 0.754 (0.290) | 0.455 (0.191) | 0.676 (2.803) | 0.008 (0.025) | 2.136 (0.141) | 2.751 (0.454) |

**Table S5.** Parameter estimates and standard deviations in brackets for full-marginalisation models across nine simulation scenarios generated from the segmented linear model with gamma error. Simulated true values for *α, β*_1_ and *β*_2_ were 0, 1 and 0.36 respectively. Estimates of *c* are expected to be near unity (see main text). A value 0.22 is expected for adult mortality from the simulated survival probabilities. The population was simulated to be on average stable i.e., *ρ* ≈ 0.

| Sample fraction | Model disp. | $\alpha$ | $\beta_1$ | $\beta_2$ | $\zeta$ | $\rho$ | $\phi$ | $c$ |
| --- | --- | --- | --- | --- | --- | --- | --- | --- |
| 0.0025 | 0.005 | 0.053 (0.049) | 0.943 (0.052) | 0.3616 (0.0791) | 0.2440 (0.0571) | -0.002 (0.0398) | 0.005 (0.5e-04) | 1.232 (0.694) |
| 0.0025 | 0.010 | 0.060 (0.060) | 0.935 (0.063) | 0.3934 (0.1015) | 0.2260 (0.0582) | 0.004 (0.0398) | 0.010 (1e-04) | 1.501 (0.803) |
| 0.0025 | 0.020 | 0.060 (0.060) | 0.940 (0.057) | 0.3966 (0.1253) | 0.2489 (0.0615) | -0.002 (0.0374) | 0.020 (2e-04) | 1.262 (0.523) |
| 0.0035 | 0.005 | 0.037 (0.044) | 0.956 (0.049) | 0.3506 (0.0804) | 0.2510 (0.0530) | -0.001 (0.0316) | 0.005 (0.5e-04) | 1.202 (0.580) |
| 0.0035 | 0.010 | 0.048 (0.047) | 0.947 (0.052) | 0.3643 (0.1008) | 0.2540 (0.0468) | 0.005 (0.0286) | 0.010 (1e-04) | 1.309 (0.441) |
| 0.0035 | 0.020 | 0.046 (0.046) | 0.947 (0.048) | 0.3531 (0.0946) | 0.2353 (0.0619) | 0.004 (0.0364) | 0.020 (2e-04) | 1.320 (0.579) |
| 0.0045 | 0.005 | 0.032 (0.039) | 0.965 (0.042) | 0.3291 (0.0678) | 0.2483 (0.0333) | -0.002 (0.0289) | 0.005 (0.6e-04) | 1.137 (0.332) |
| 0.0045 | 0.010 | 0.034 (0.036) | 0.960 (0.037) | 0.3335 (0.0686) | 0.2409 (0.0461) | 0.008 (0.0233) | 0.010 (1e-04) | 1.275 (0.473) |
| 0.0045 | 0.020 | 0.029 (0.035) | 0.963 (0.043) | 0.3461 (0.1084) | 0.2380 (0.0347) | 0.010 (0.0226) | 0.020 (2e-04) | 1.375 (0.414) |

## Supplementary Figures

**Figure S1.**
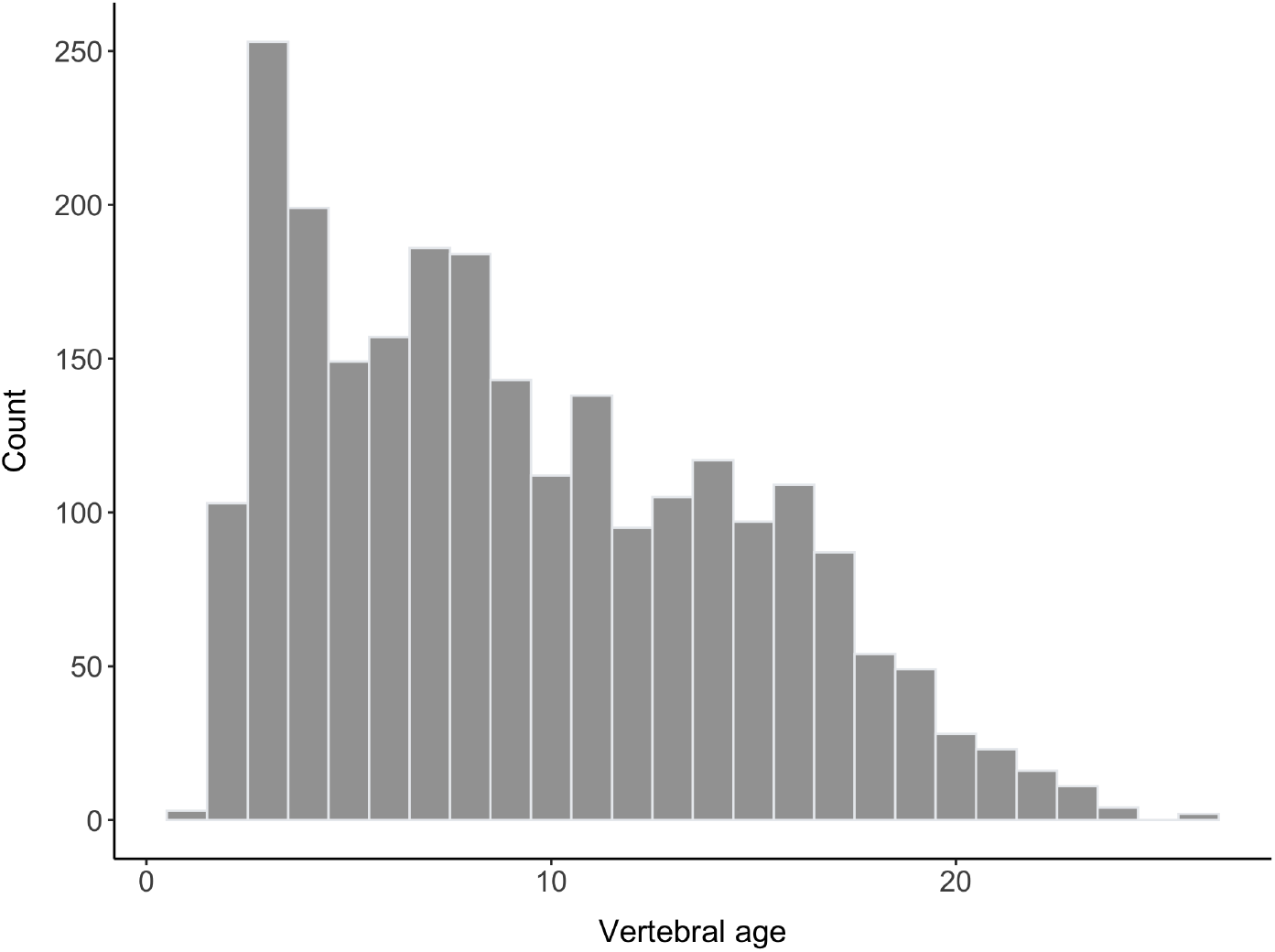
Distribution of available vertebral band counts from 2,424 school shark samples.

**Figure S2.**
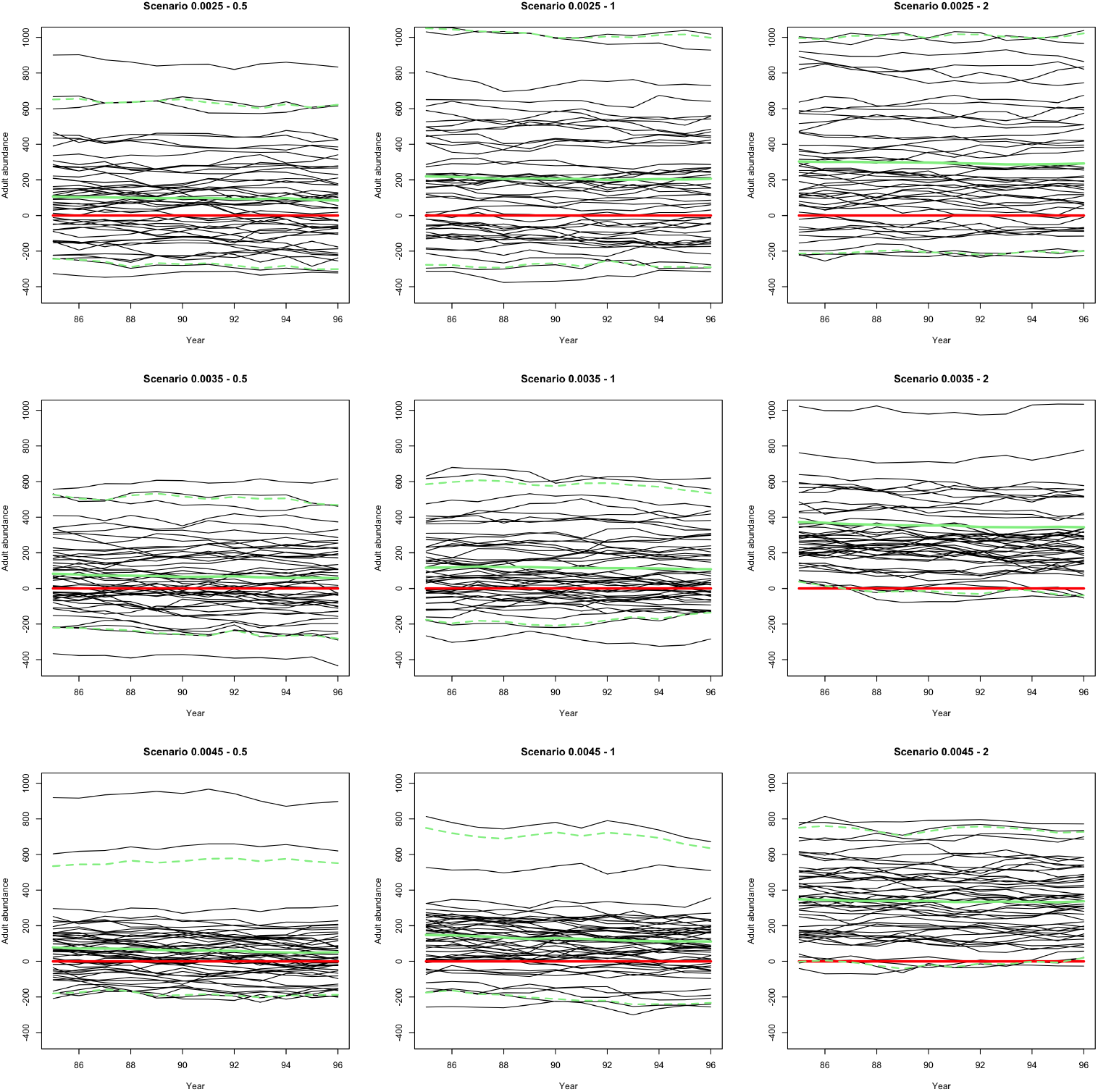
Time-series summary of estimated adult abundance differences (from simulated truth) from pairwise difference model from normal linear model simulation scenario. Each panel shows the results from a simulation scenario with increasing sample size from top to bottom and increasing simulated error variance from left to right. Panel headings describe the sample proportion and assumed standard deviation for the normal error. Red line shows the zero line. Solid green line is the mean over the simulation replicates and the dashed green lines the 95% within-year quantiles over replicates. Lines are straight because the pairwise difference model averages the demographic model over time.

**Figure S3.**
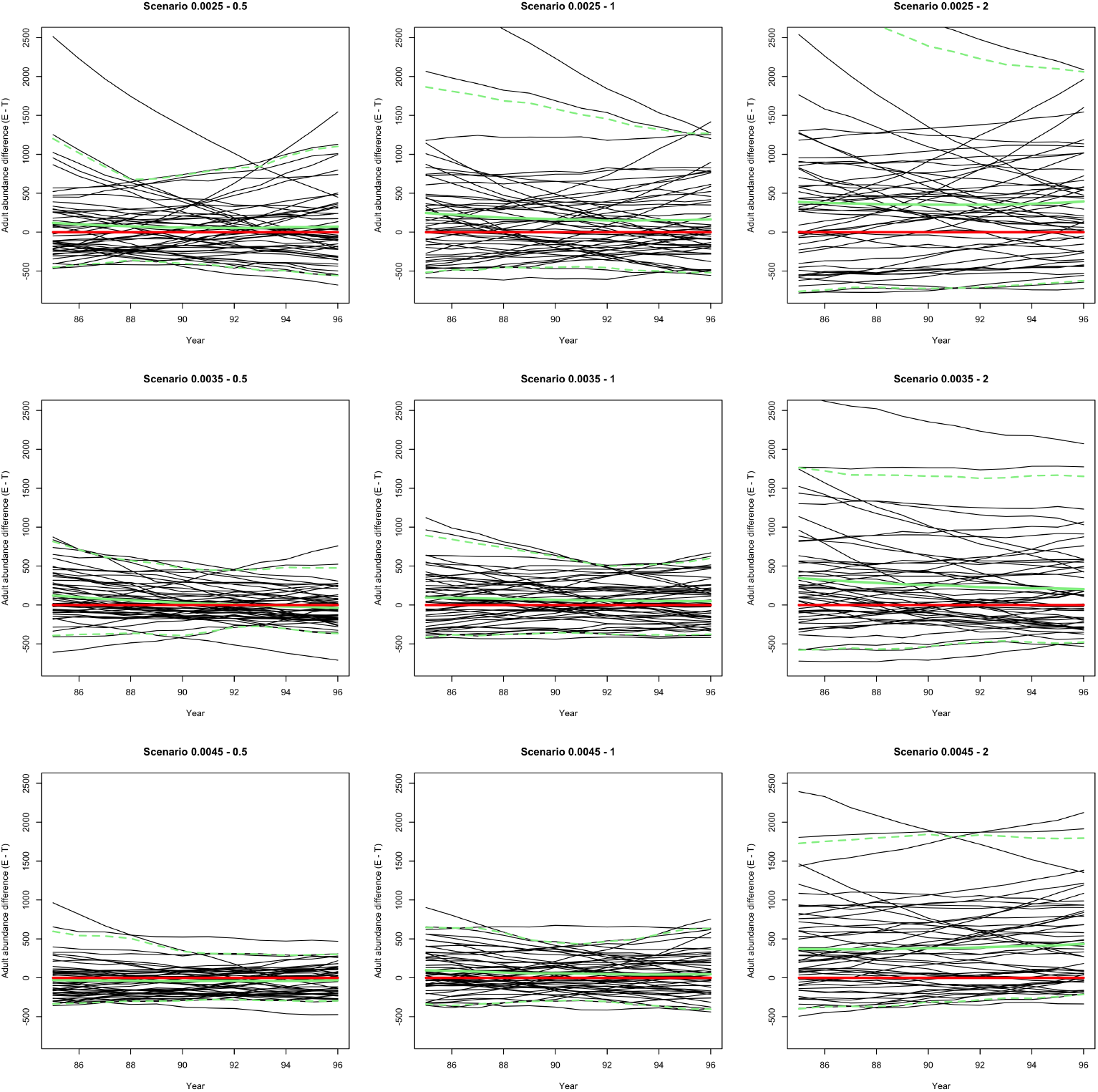
Time-series summary of estimated adult abundance differences (from simulated truth) from full-marginalisation model from normal linear model simulation scenario. Each panel shows the results from a simulation scenario with increasing sample size from top to bottom and increasing simulated error variance from left to right. Panel headings describe the sample proportion and assumed standard deviation for the normal error. Red line shows the zero line. Solid green line is the mean over the simulation replicates and the dashed green lines the 95% within-year quantiles over replicates.

**Figure S4.**
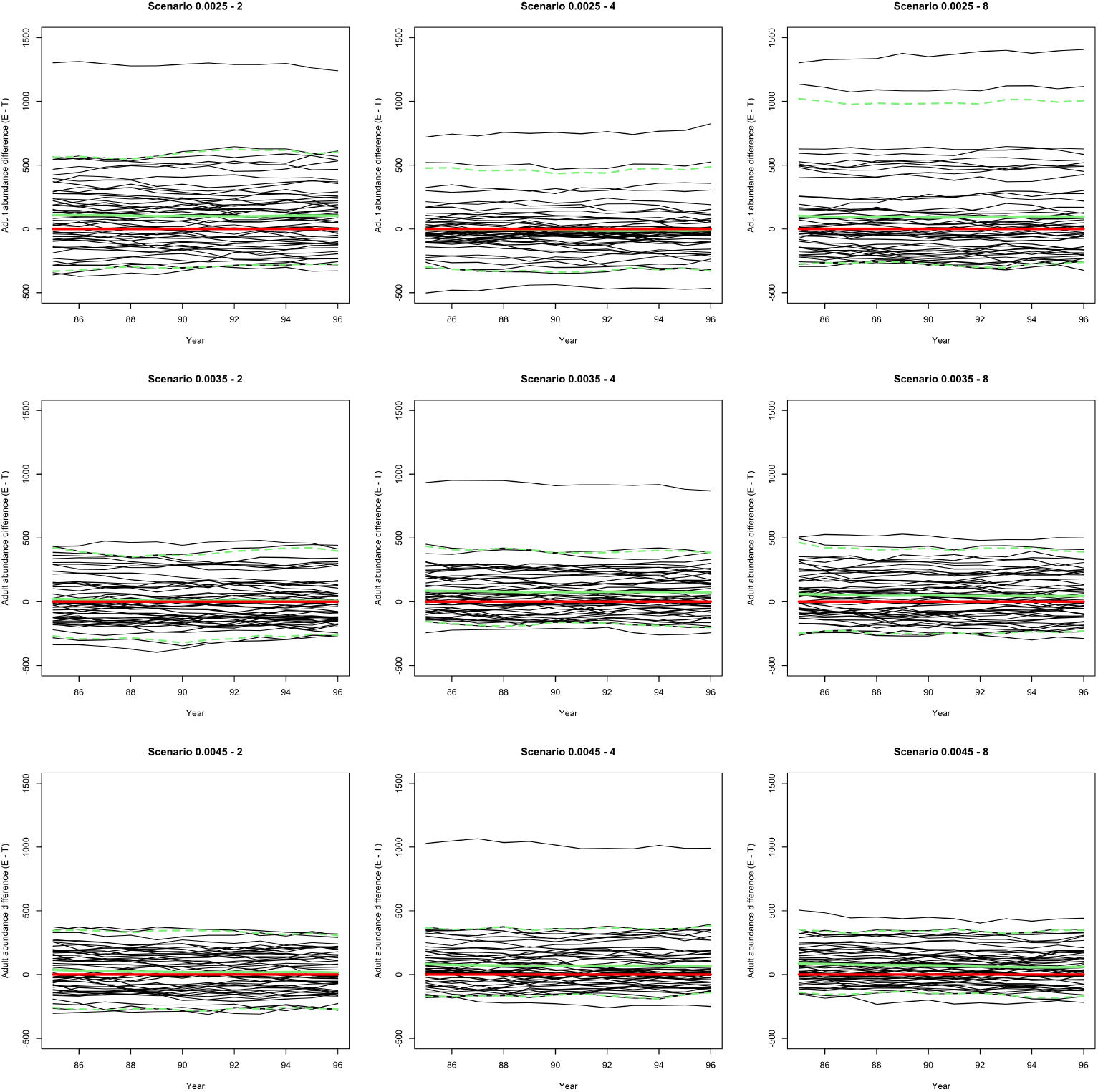
Time-series summary of estimated adult abundance differences (from simulated truth) from pairwise difference model from the scaled normal linear model simulation scenario. Each panel shows the results from a simulation scenario with increasing sample size from top to bottom and increasing simulated error variance from left to right. Panel headings describe the sample proportion and assumed standard deviation for the normal error. Red line shows the zero line. Solid green line is the mean over the simulation replicates and the dashed green lines the 95% within-year quantiles over replicates.

**Figure S5.**
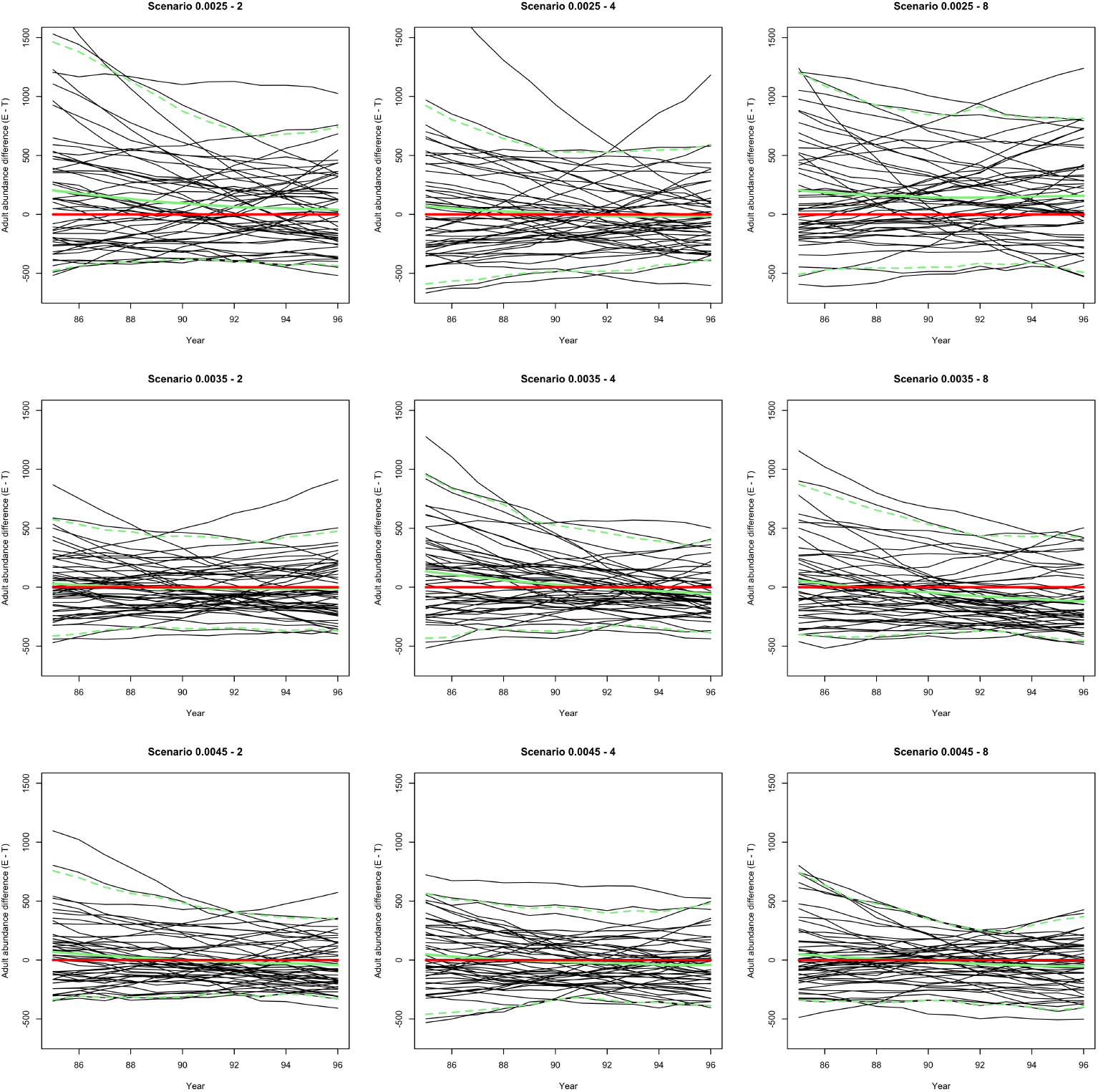
Time-series summary of estimated adult abundance differences (from simulated truth) from full-marginalisation model from the scaled normal linear model simulation scenario. Each panel shows the results from a simulation scenario with increasing sample size from top to bottom and increasing simulated error variance from left to right. Panel headings describe the sample proportion and assumed standard deviation for the normal error. Red line shows the zero line. Solid green line is the mean over the simulation replicates and the dashed green lines the 95% within-year quantiles over replicates.

**Figure S6.**
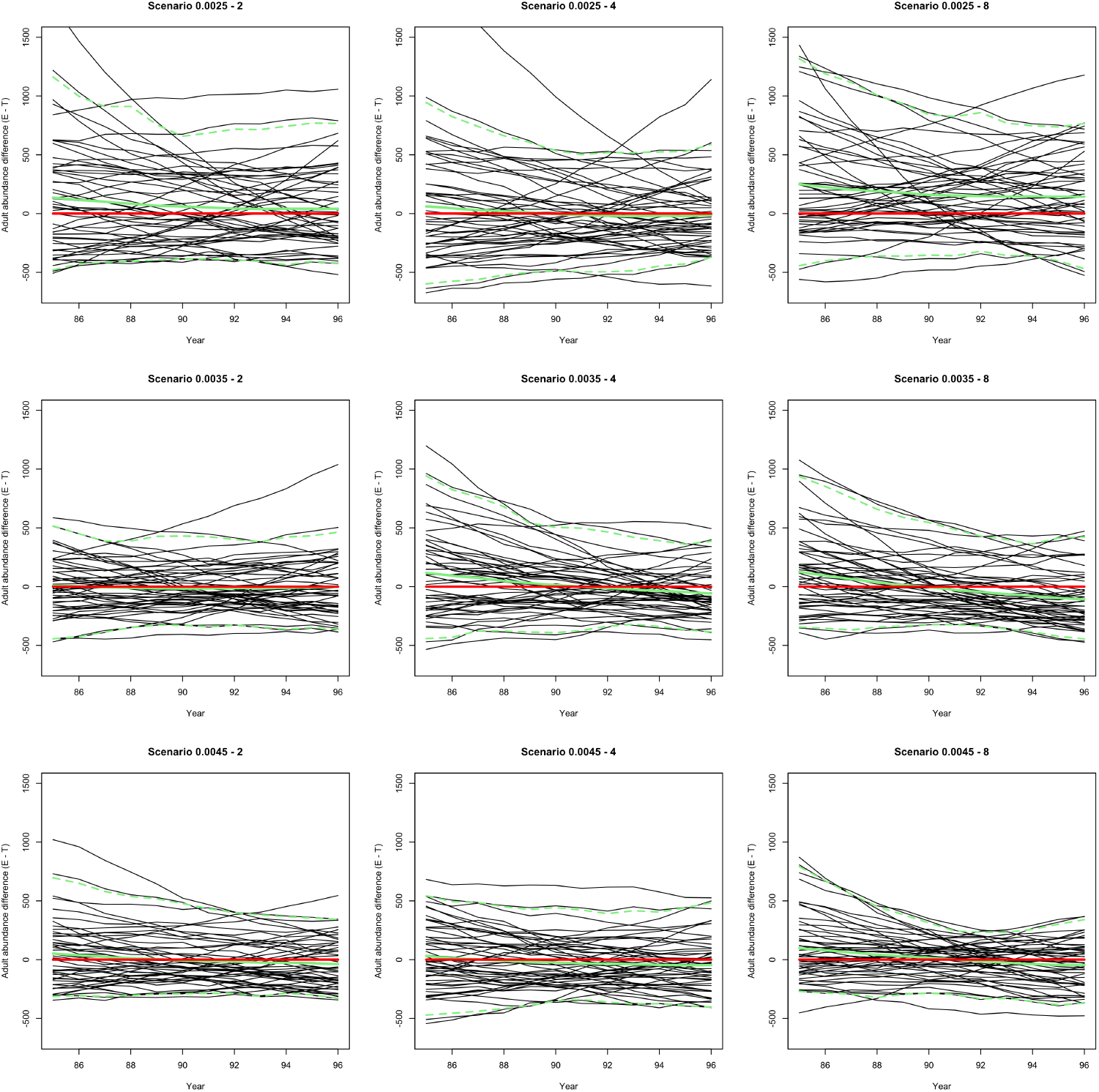
Time-series summary of estimated adult abundance differences (from simulated truth) from full-marginalisation model from the scaled normal linear model simulation scenario *and assumed prior means* for *σ* deviating from the truth. For simulated models with S.D.s of 2 and 4, the prior mean is set to 50% more and for 8 it is set to half that value. Each panel shows the results from a simulation scenario with increasing sample size from top to bottom and increasing simulated error variance from left to right. Panel headings describe the sample proportion and assumed standard deviation for the normal error. Red line shows the zero line. Solid green line is the mean over the simulation replicates and the dashed green lines the 95% within-year quantiles over replicates.

**Figure S7.**
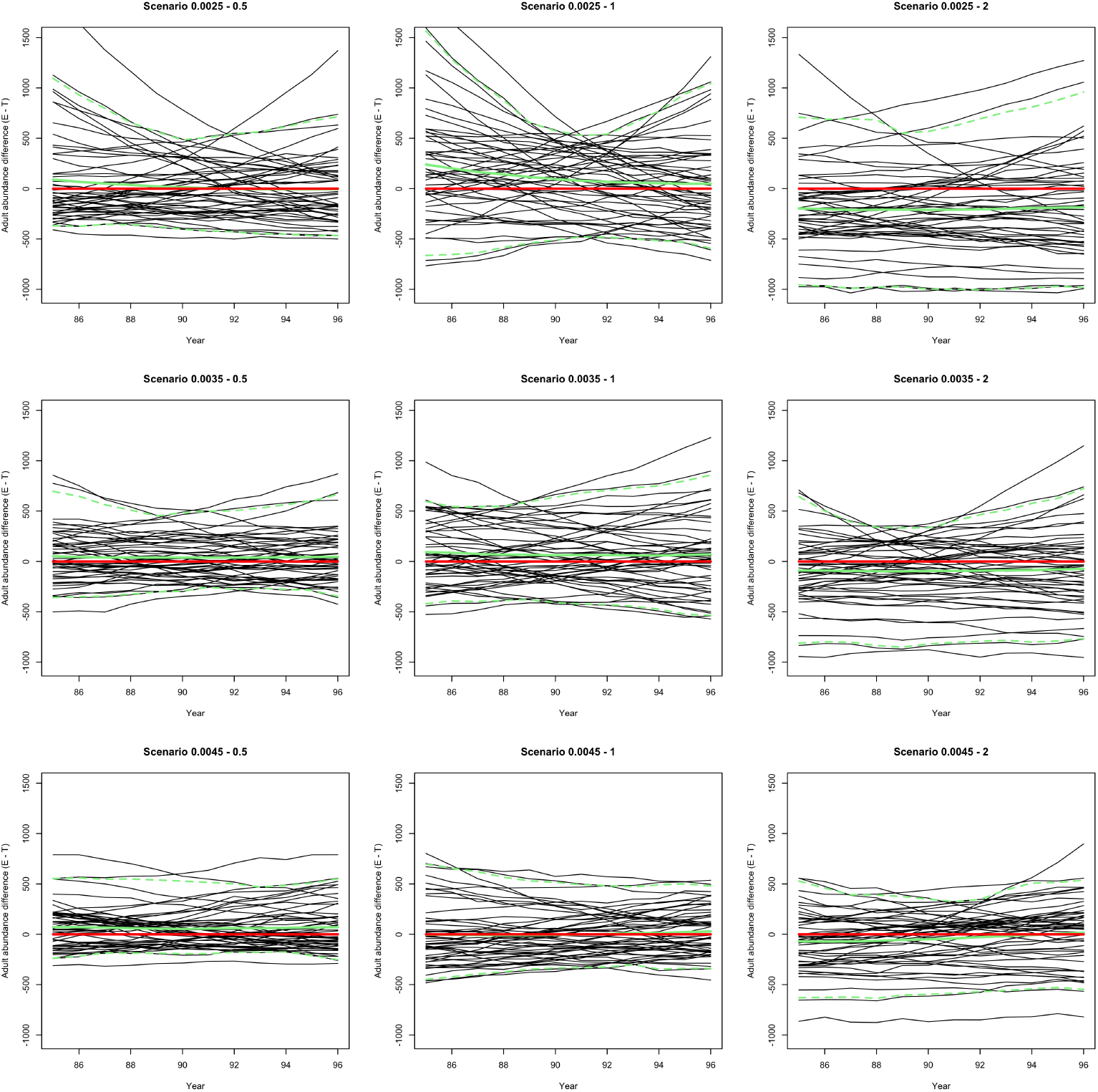
Time-series summary of estimated adult abundance differences (from simulated truth) from full-marginalisation model from the segmented linear model with normal error simulation scenario. Each panel shows the results from a simulation scenario with increasing sample size from top to bottom and increasing simulated error variance from left to right. Panel headings describe the sample proportion and assumed standard deviation for the normal error. Red line shows the zero line. Solid green line is the mean over the simulation replicates and the dashed green lines the 95% within-year quantiles over replicates.

**Figure S8.**
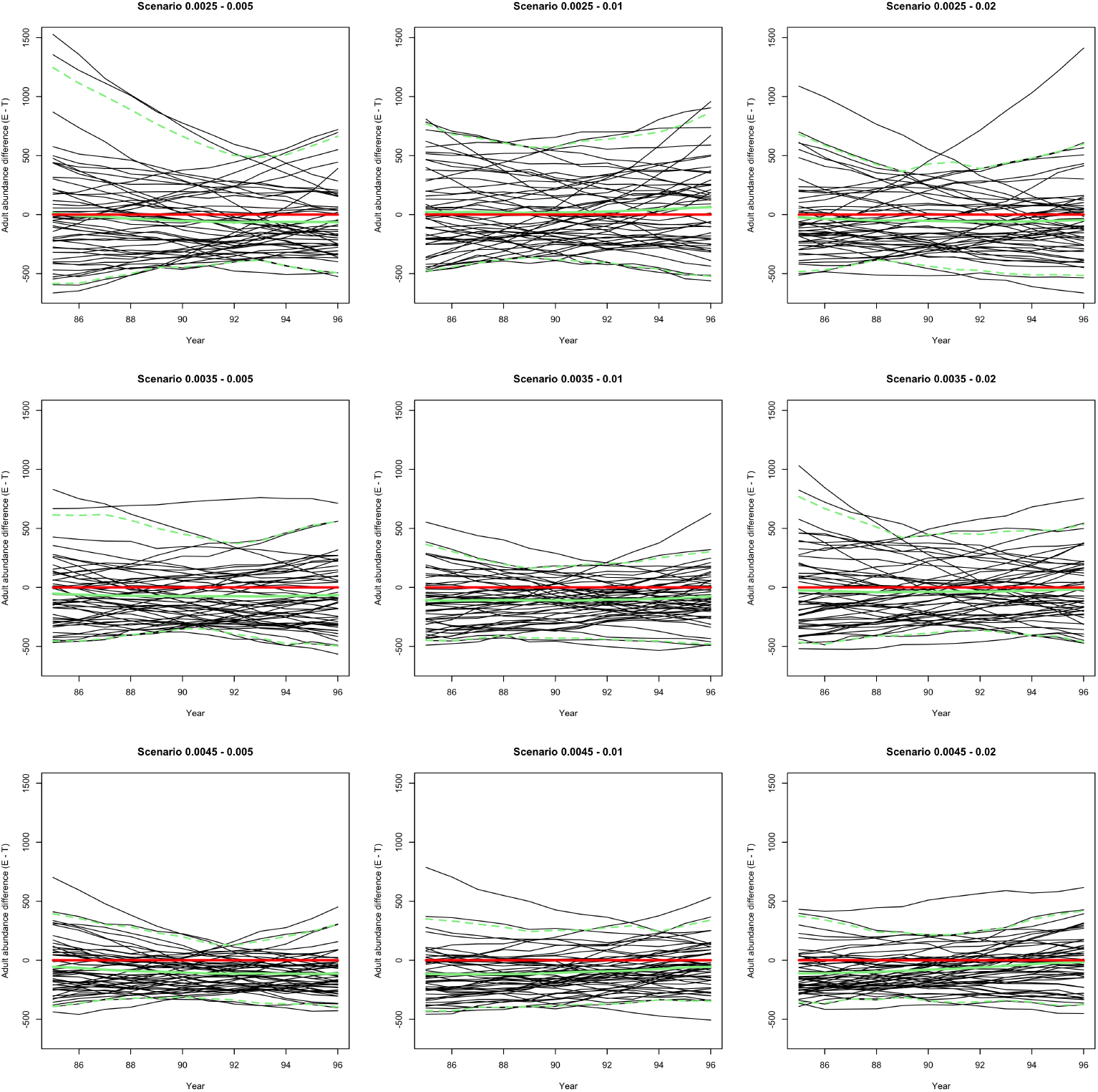
Time-series summary of estimated adult abundance differences (from simulated truth) from full-marginalisation model from the segmented linear model with gamma error simulation scenario. Each panel shows the results from a simulation scenario with increasing sample size from top to bottom and increasing simulated error variance from left to right. Panel headings describe the sample proportion and assumed standard deviation for the normal error. Red line shows the zero line. Solid green line is the mean over the simulation replicates and the dashed green lines the 95% within-year quantiles over replicates.

